# Transposon invasion of primate genomes shaped human inflammatory enhancers and susceptibility to inflammatory diseases

**DOI:** 10.1101/2025.01.21.632732

**Authors:** Mengliang Ye, Maxime Rotival, Sebastian Amigorena, Elina Zueva

## Abstract

Human immune inflammatory response reflects the evolutionary adaptation of immune-cell regulatory elements^1^, where recent mutations can control both pathogen defence and susceptibility to chronic inflammatory and autoimmune diseases^2–4^. The impact of the deeper evolutionary history of these elements within primate genomes on human inflammatory responses remains poorly understood. Evolutionary young transposons have uniquely reshaped primate genomes^5^ and spread novel *cis*-regulatory sequences^6^. To understand how these events influenced human inflammation, we traced sequence changes in annotated human immune-cell enhancers back to macaque. We show that Alu elements and endogenous retroviruses dispersed motifs for the inflammation-related NF-κB and IRF1, redefining their binding patterns and contributing most prominently to great ape-specific binding sites. After the human-macaque split, many of these motifs shifted toward higher predicted binding affinity. In humans, population genetics analyses reveal that positive selection favors alleles, often Alu-derived, that increase enhancer affinity toward NF-κB. Enhancers containing Alu elements are more likely to undergo positive selection at the locus level, particularly when associated with chronic inflammatory diseases. As the most mutable enhancer sequences, Alus harbor disproportionately high numbers of single nucleotide polymorphisms and significantly contribute to selected alleles, some associated with chronic inflammatory diseases. We propose that the invasion of primate-specific transposons has created unique opportunities to adapt inflammatory responses in rapidly evolving great apes, with ancestral Alus continuing to influence evolutionary potential in humans.

## Main Text

Inflammatory immune responses are vital for survival, acting as a defense mechanism against various threats. However, inflammation is also involved in nearly all modern diseases^7,8^. The genetic roots of this paradox are embedded in the human evolutionary history. For instance, mutations selected to bolster immune defense in past environments can, under modern conditions, heighten the risk of chronic inflammation^2–4,9^. Evolutionary events preceding human history may have laid a more ancient foundation for contemporary inflammatory responses. A recent study shows that great apes, including humans, exhibit a more robust and broad early transcriptional response to immune stimulation compared to monkeys^10^. This response is enriched in inflammatory pathways^10^, suggesting rapid regulatory evolution of inflammation in hominids. While this adaptation may protect population fitness in primates with increased body size and delayed reproductive maturity, it may also predispose to chronic inflammatory diseases.

Inflammatory responses are under the control of transcriptional enhancers that often harbor risk alleles and are recognized as crucial elements of evolutionary adaptability^1^. Enhancers evolve through mutations and expansion of DNA sequences, altering transcription factor binding sites (TFBS) and, thereby, gene expression^11–14^. During evolution, transposable elements (TEs) represent one of the major sources of novel DNA^12,15^ and provide not only raw DNA material but also regulatory motifs, facilitating enhancer evolution^16^. Though most TEs are dormant today, they once spread to occupy nearly half of the mammalian genomes^5^. Based on their transposition mechanism - either through DNA or RNA intermediates - TEs are classified into DNA transposons and the more prevalent retrotransposons, which include endogenous retroviruses (ERVs), long interspersed nuclear elements (LINEs), and short interspersed nuclear elements (SINEs, including Alus in primates)^17^, all further branching into smaller groups and subfamilies^18^.

Throughout primate evolution, periodic invasions by “primate-specific” TEs have profoundly reshaped genomes and regulatory networks^6,12,19,20^. The link between these TEs and immunity has been recognized previously. For example, retrovirus Mer41B, through interferon response motifs, contributes to the IFN-I responses associated with immunity and inflammation^21^. Alus are particularly biased towards immune-cell regulatory elements^22^ and, along with other primate-specific TEs, play a significant role in the formation of novel enhancers^6^. Even in non-immune tissues like the liver, active enhancers harboring primate-specific transposons are located near genes linked to immunity and inflammation^6^.

Here, we used comparative genomics and population genetics to systematically explore how primate-specific transposable elements (pTEs) have shaped the evolution of human immune-cell enhancers across distinct evolutionary time scales. By reconstructing the history of sequence divergence, pTE acquisition, and the emergence of proinflammatory TFBS following the human-macaque split, we show that pTEs introduced a substantial number of motifs for inflammation-related transcription factors. Integrating large-scale ChIP-seq datasets, we reveal that these pTE-derived motifs actively promote the *in vivo* binding of the corresponding proinflammatory NF-κB and IRF1 proteins. The significance of pTE-derived binding sites has grown over evolutionary time as they became key contributors to functional great-ape-specific TFBS. Furthermore, NF-κB1 and IRF1 motifs derived from Alu elements and primate ERVs, respectively, exhibit signatures of fine-tuning in primates and humans, with a consistent trend toward increased predicted binding affinity. We emphasize the important role of Alus as highly mutable elements and hotspots of positive selection in humans, serving as a major source of NF-κB1 motifs under selection. Our findings highlight the profound impact of ancestral pTEs on the evolution of human inflammatory responses and reveal a vast reservoir for future adaptations.

## Results

### Primate-specific TEs are enriched in rapidly diverging immune-cell enhancers

To investigate the role of pTEs in the regulatory evolution of human inflammatory response, we first compiled a comprehensive list of putative human immune-cell enhancers. We extracted annotations for various lymphoid and myeloid populations from Enhancer Atlas 2.0^23^ and for lymphoblastoid cell lines from *Garcia-Perez et al.*^24^. To select robustly active enhancers, we integrated published ATAC-sequencing across immune cell types^25^. This yielded 60,332 regions broadly shared by distinct immune cell types (Extended Data Fig.1a) and frequently containing multiple ATAC peaks (Extended Data Fig.1b). These regions, characterized by dynamic chromatin opening with fluid boundaries between cell types, have a median length of 3kb after merging overlapped coordinates (Extended Data Fig.1c). Given their promiscuous profile, we refer to them collectively as “immune-cell enhancers.”

To trace DNA sequence evolution within these human immune-cell enhancers back to macaque, we converted their coordinates to high-quality genome assemblies of rhesus macaque (RheMac10) and chimpanzee (PanTro6), representing our closest monkey and great ape relatives, which diverged approximately 30-35 and 6-12 million years ago, respectively^26^. Since human and chimpanzee genomes differ by only a few percent^27^, we defined orthologs as enhancers with a minimum of 97% alignability between species. Using the *UCSC LiftOver* converter, we classified enhancers into three groups based on their alignability under these conditions: i) three-species orthologs (30,058), ii) human/chimpanzee orthologs (25,479) carrying human-chimp sequence variations, and iii) regions with human-specific variations (4,795) (Fig.1a). Based on the evolutionary timing of their sequence divergence, these groups were defined as “static,” “intermediate,” and “rapid,” with the latter two collectively referred to as “dynamic” regions (Fig.1a). While dynamic regions were not alignable to the macaque genome at a given threshold, lowering it to 50% alignability identified quasi-orthologs for 96% of them, indicating that the alteration of ancestral sequences is more prevalent than the emergence of enhancers from entirely novel DNA.

**Fig. 1.**
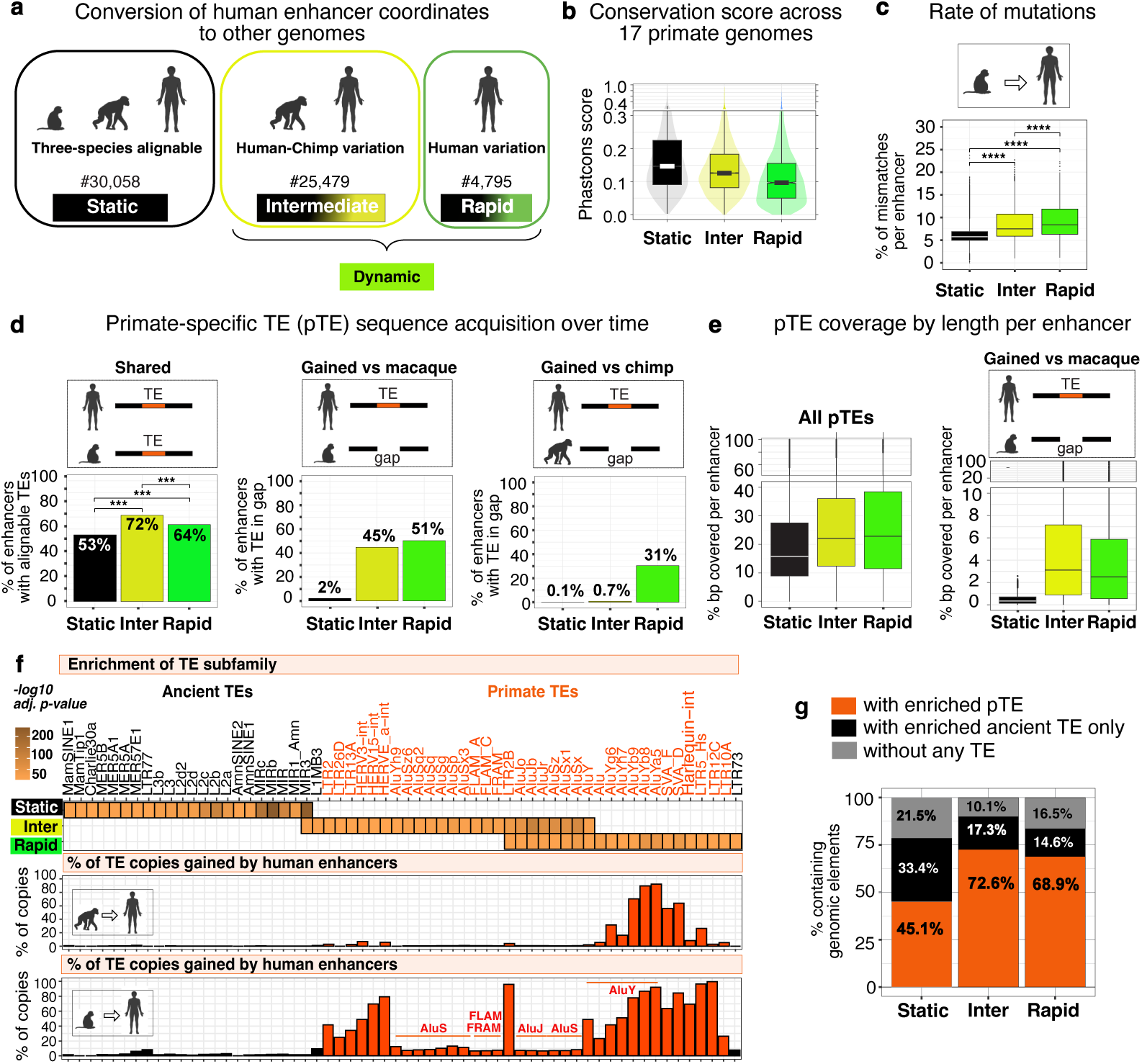
Evolution of sequences in regions corresponding to human immune-cell enhancers since the human-macaque split. **a,** Stratification of human enhancer regions by evolutionary age based on their alignment to the macaque and chimpanzee genomes. The number of enhancers in each group is indicated. **b,** *PhastCons* conservation scores for enhancer groups across 17 primate genomes. **c,** Percentage of small-scale mutations in enhancers based on *BLASTn*. **d,** Proportion of enhancers overlapping with primate-specific TEs (pTEs), either present in the same genomic location in macaque (shared) or corresponding to genomic gaps (gained) in macaque (middle panel) or chimpanzee (right panel). *\*\*\*P ≤ 0.001*. **e,** Distribution of the percent of enhancer length covered by pTE sequences, considering either all pTE (left) or the subset of pTEs that are absent from the macaque genome (right). **f, Upper panel:** Top TE subfamilies enriched in distinct enhancer groups as determined by the hypergeometric test (*adj. P < 10^−7^ | FC ≥ 2 & nCount ≥ 10*), while not enriched in random sets of genomic regions matched to the corresponding enhancer group by number and size distribution, and shuffled 1000 times. **Middle and lower panels:** Proportion of the enhancer pTE copies gained since the split from chimpanzee (middle panel) or macaque (lower panel), **g,** Proportion of enhancers either overlapping with pTE or ancient TE (without pTE overlap) or not containing TE sequence.

We used *Phastcons* to measure sequence conservation across 17 primate clades, revealing that dynamic regions are less conserved than static regions over a broader evolutionary scale (Fig.1b). We then measured mutation rates without accounting for larger insertions and deletions. Using BLASTn, we quantified nucleotide mismatches between human and macaque orthologs (for static regions) and between human and macaque quasi-orthologs (for dynamic regions). Dynamic regions exhibited a higher frequency of mutations per enhancer compared to static regions (Fig.1c). Together, these results suggest that, compared to static regions, dynamic ones have evolved under lower genetic constraint, i.e., weaker purifying selection, which permits a higher TE frequency^28^. To measure pTE content in distinct enhancer groups, we annotated TEs using RepeatMasker (www.repeatmasker.org) and distinguished primate-specific biotypes using clade information from the TEanalysis tool (https://github.com/4ureliek/Teanalysis). Focusing on the timing of pTE acquisition following the human-macaque split, we classified pTE sequences into two categories: those inherited from the human-macaque common ancestor, identified by their conserved genomic locations between human and macaque (“shared” pTEs) and those acquired later in evolution, identified by their presence within enhancers in humans but corresponding to genomic gaps in macaque or chimpanzee (“gained” pTEs). The “gained” pTEs could have originated from the reactivation of pre-inserted pTE species or the invasion of newly emerging ones. Although likely a minority, some evolutionary gains may have arisen from the loss of the original copy due to genome re-shuffling during speciation.

pTE content analysis revealed that, while a high proportion of enhancers in each of the three groups contain “shared” pTE sequences (Fig.1d, left panel), dynamic enhancers are more likely to overlap with such pTEs compared to static enhancers (72% and 64% vs. 53%). They were also more likely to acquire pTEs following the split from macaque, with nearly half containing pTE sequence absent in macaque, compared to only 2% of static enhancers (Fig.1d, middle panel), and with rapid regions accounting for the majority of further human-specific gains (31%) (Fig.1d, right panel). Furthermore, dynamic regions show a greater median length covered by pTE sequences (∼20% in bp per enhancer) compared to static regions (∼16%) (Fig.1e, left panel). Among these pTEs, those acquired after the human-macaque split contribute a median of 2–3% to the length of dynamic enhancers, compared to only 0.2–0.3% in static regions (Fig.1e, right panel). Together, these patterns correspond to the low purifying selection identified above in dynamic regions, as this facilitated pTE accumulation.

To identify TE subfamilies specifically enriched in distinct enhancer groups, we compared their abundance within enhancers to their genomic frequencies and 1,000 times shuffled random genomic regions matched in number and size distribution to each enhancer group. Ancient TE subfamilies were significantly enriched in static enhancers, while pTE subfamilies were predominantly enriched in dynamic enhancers (Fig.1f, upper panel). Of the 485 pTE subfamilies in the human RepeatMasker, 55 were enriched in dynamic enhancer groups. Over 70% of dynamic regions overlapped with pTEs from these enriched subfamilies, compared to less than 50% of static regions (Fig.1g). We traced the acquisition of the enriched TE species over time by mapping their human coordinates to the corresponding genomic gaps in macaque and chimpanzee. As expected, most ancient TE copies were gained before the human-macaque split, as their locations are conserved (Fig.1f, middle and lower panels). For pTEs, acquisition timing corresponded to their known evolutionary birth age. For example, up to 80% of the youngest AluY elements map to genomic gaps in chimpanzee and macaque, representing human-specific gains (Fig.1f, middle and lower panels). In contrast, older AluJ, which colonized early primate genomes, and AluS, which newly emerged in monkeys^29^, are primarily shared between humans and chimpanzees. However, 10–15% of both AluJ and AluS copies are absent in macaque, suggesting their reactivation in the common human-chimpanzee ancestor.

Alu elements, 300-base retrotransposons derived from the 7Sl gene^30^, are the most abundant transposons in the human genome^29^. In immune-cell enhancers analyzed here, Alus account for 98% of all enriched pTE copies, compared to 67% in the broader genome (Extended Data Fig.2a). Alus are known to carry enhancer-specific histone marks at nucleosomes^31^. Meanwhile, less abundant ERVs are rich in regulatory sequences^16^. We examined the contribution of these two pTE biotypes to the enhancer’s nucleosome-free open chromatin that harbors key hotspots for transcription factor binding. Analysis of ATAC-seq peaks from immune cell subsets revealed that Alus contribute up to 1.8% and primate-specific ERVs (pERVs) up to 4% of peak summits within DNA shared with macaque (Extended Data Fig.2b). This contribution increased sharply in evolutionarily novel DNA absent from the macaque genome, with pERVs and Alus each generating a significant proportion of open chromatin, accounting for up to 16% and 20%, respectively (Extended Data Fig.2b and Extended Data Fig.2c for an example). These results demonstrate that pTE sequences gained after the human-macaque split were rapidly co-opted for regulatory purposes, with this process being particularly efficient for Alu elements, whose contribution to open chromatin increased up to 11-fold compared to the earlier-inserted counterparts (Extended Data Fig.2b, right panel).

In summary, we identified human immune-cell enhancer regions that were particularly prone to accumulating pTE sequences after the human-macaque split, likely due to low purifying selection. While pTEs generated regulatory hotspots of open chromatin within enhancers, their impact was especially pronounced during the latest stages of primate evolution.

### Dynamic regions are primarily enriched for inflammation-related TFBS

To investigate the evolution of inflammation-related transcription factor (TF) binding sites across enhancer groups, we first systematically scanned these regions for over-represented TFBS using the HOMER collection of ChIP-seq-derived motifs. Compared to genomic background, static regions were enriched for a wide array of TFBS, including for the inflammation-related IRF family proteins, which mediate interferon I response during viral infections and inflammation^32^, and NF-κB, critical for initiating and resolving inflammatory responses^33^ (Fig.2a). Notably, NF-κB is a multiprotein family including NF-κB1, NF-κB2, RELA, RELB, and RELC, all of which bind to variations of the same consensus sequence^33^. Additional enriched TFBS in static regions included those for NF-E2-related factors, antagonists of NF-κB^34^, and JUN, ETS, and RUNX family proteins, playing diverse roles in immunity^35,36^. Enrichments were also observed for zinc finger proteins and immune cell identity factors. In contrast, dynamic regions were enriched for a more limited set of predominantly inflammation-related TFBS (Fig.2a).

**Fig. 2.**
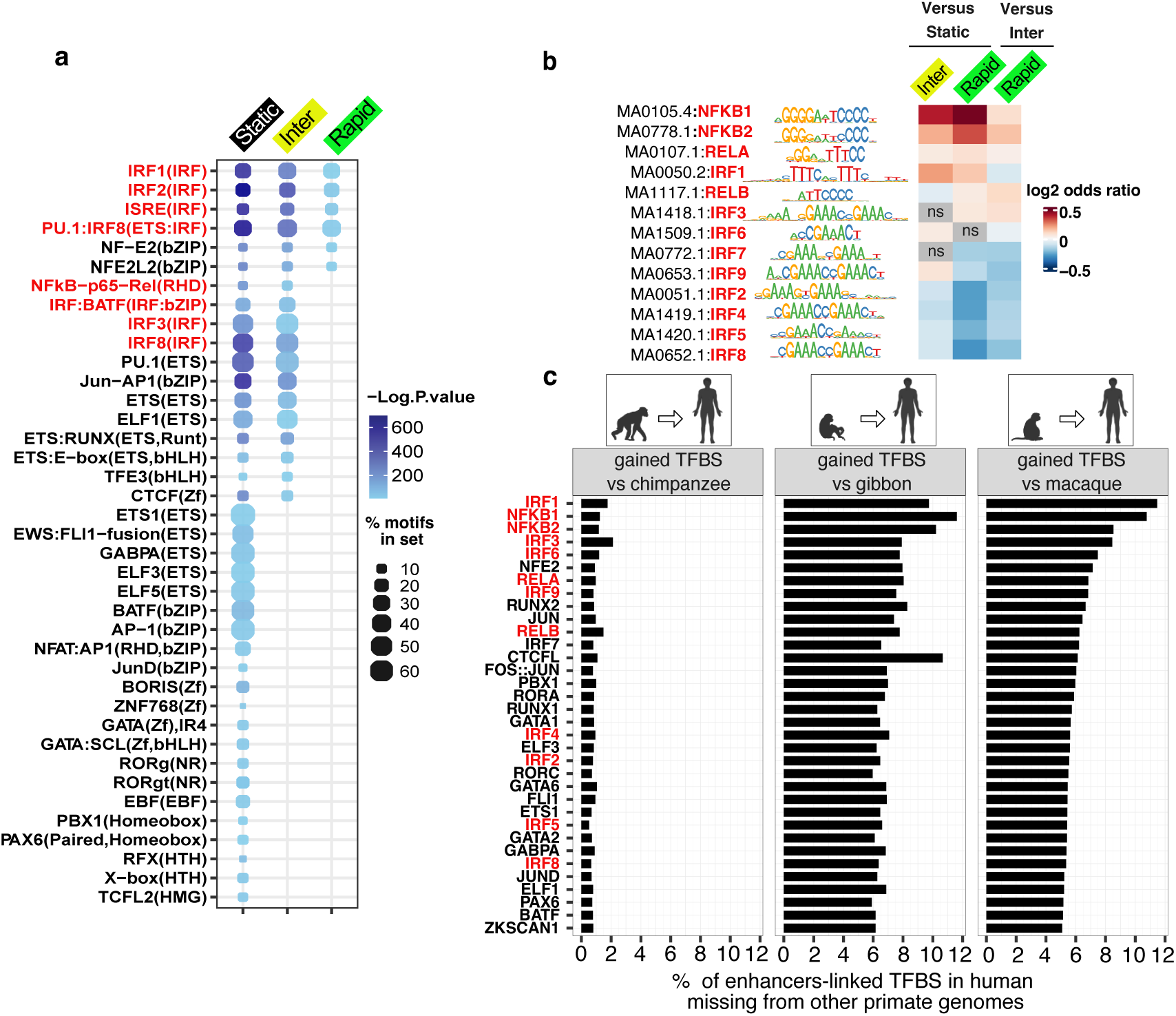
Evolution of the enhancer regulatory lexicon. **a,** TFBS from Homer collection of Chip-seq motifs that are significantly enriched in distinct enhancer groups *(P <0.01, % of motifs in set ≥5, Fold change ≥10)*. **b,** Motif enrichment for NF-κB and IRF family proteins in intermediate enhancers compared to static enhancers (used as background), and in rapid enhancers compared to either static or intermediate enhancers (used as background). **c,** Proportion of TFBS in human immune-cell enhancers that match gaps in the genomes of macaque (RheMac10), gibbon (NomLeu3), or chimpanzee (PanTro6), relative to the total number of the corresponding motifs in enhancers. **Panels a-c**: Inflammation-related motifs are highlighted in red.

To understand the reasons behind these distinct patterns, we focused on motifs for inflammation-related IRF and NF-κB factors using the JASPAR2022 collection, comparing the two dynamic groups to static regions and the rapid group to the intermediate group. We found that dynamic regions were significantly more enriched for NF-κB (NF-κB1, NF-κB2, and RELA) and IRF1 motifs compared to static regions (*adj. P-value* <10^-10^). Furthermore, intermediate regions were more enriched for IRF1, IRF6, and IRF9 motifs, while rapid ones were more enriched for RELB and IRF3 motifs (Fig.2b). This suggests a larger pool of proinflammatory motifs in dynamic regions compared to static regions.

To trace the evolutionary emergence of additional TFBS, we focused on TF families enriched in enhancer groups. Human JASPAR2022 USCS coordinates of their respective TFBS in enhancers were intersected with the corresponding genomic gaps in chimpanzee and macaque, with gibbon added as a lesser ape to identify great-ape-specific gains. Quantifying the proportions of examined TFBS types that were gained (map to the corresponding gaps) since the human-macaque split revealed that most of them are shared between human and chimpanzee, with only ∼1% being human-specific (Fig.2c, left panel). In contrast, 6% to 11% are absent from the gibbon and macaque genomes (Fig.2c). Given that most *de novo* TFBS are absent from both genomes, we broadly classified these sites as “great-ape-specific.” Among these, motifs for proinflammatory factors IRF1 and NF-κB1 showed the most rapid gains, evidenced by the highest proportion of *de novo* motifs relative to their respective total pools (Fig.2c). These *de novo* TFBS nearly exclusively emerged within dynamic regions (Extended Data Fig.3), now accounting for thousands of potential binding sites.

To further understand the functional roles of enhancer groups, we linked them to immune-response-related genes from^10^ using the activity-by-contact (ABC) enhancer-gene interaction maps across immune populations^37^. This revealed a significant overlap in gene targets between distinct enhancer groups (Extended Data Fig.4), implying regulation of the same functional pathways.

In conclusion, inflammation-related TFBS present within human immune-cell enhancers but absent in macaque, were primarily gained during the evolution of the human-chimpanzee common ancestor. These TFBS predominantly emerged within dynamic regions, where IRF1 and NF-κB motifs expanded more rapidly than other types of TFBS. Our data suggest distinct *modus operandi* among enhancer groups, with static regions regulating conserved aspects of immunity, whereas dynamic regions optimizing inflammatory responses and playing a particularly important during the evolution of the human-chimpanzee common ancestor.

### pTEs disseminated inflammation-related TFBS, reshaping transcription factor binding in immune-cell enhancers

To assess the role of pTEs in inserting TF binding sites, we overlapped selected TFBS coordinates from the human JASPAR2022 tracks with pTE enhancer locations. We retained the previous classification of TFBS as “shared” (with conserved human-macaque locations) and “great-ape-specific” (matching genomic gaps in macaque). The analysis revealed that pTEs made significant contributions to “shared” TFBS, accounting for 10% to 50% of the sites, depending on the TF type (Fig.3a, left panel). At the same time, their contribution to “great-ape-specific” TFBS is markedly higher, with pTEs in some cases serving as a critical source, particularly for NF-κB-related transcription factors (Fig.3a, right panel). For example, Alu elements contribute over 70% of “great-ape-specific” NF-κB1 motifs and more than 40% of NF-κB2 motifs. Most of these Alu elements (90%) perfectly match genomic gaps in macaques (with a minimum 97% overlap), suggesting that new Alu insertions were the primary drivers of NF-κB motif expansion during the evolution of the common human-chimpanzee ancestor. Noteworthy, Alus are not the most frequent sequences that underlie TFBS-corresponding gaps. Most other “great-ape-specific” TFBS originate from the non-Alu DNA (Fig.3a, right panel).

**Fig. 3.**
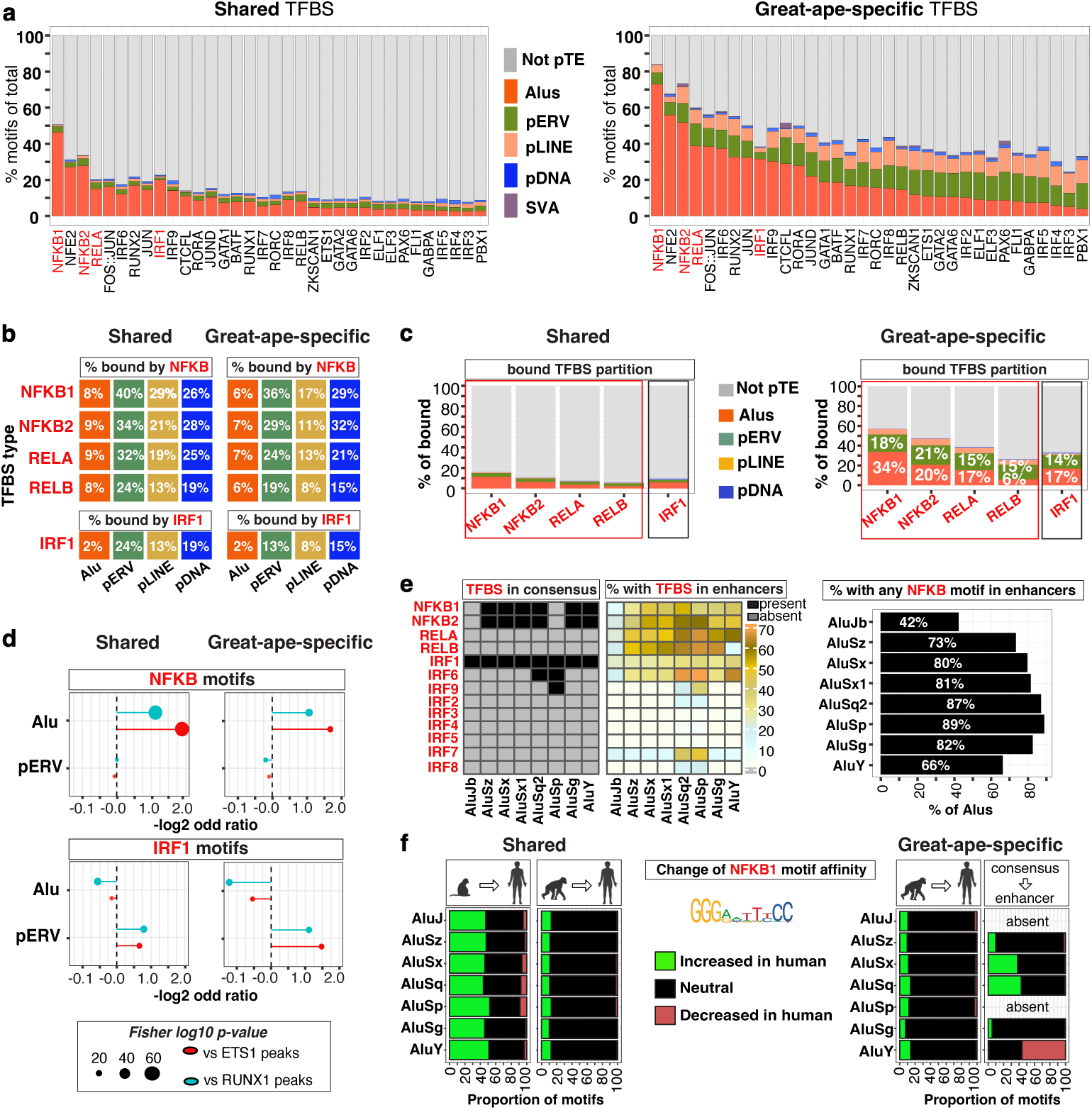
Evolution and functionality of the inflammation-related pTE-derived TFBS. **a,** Proportional contribution of distinct sequence types to individual TFBS in enhancers, with pTE-derived TFBS highlighted in color. **Left:** TFBS shared with macaque. **Right:** great-ape-specific TFBS (*fully contained within genomic gaps in RheMac10 compared to human genome GRCh38-hg38*). **b,** Proportion of TFBS derived from distinct pTE types and bound by their corresponding proteins in ChIP-seq. **Left:** TFBS shared with macaque. **Right:** great-ape-specific TFBS**. c,** Proportional contribution of distinct sequence types to shared (left panel) or great-ape-specific (right panel) TFBS bound in enhancers by their corresponding proteins, as defined by ChIP-seq. pTEs are highlighted in color **d,** Enrichment or depletion of Alu- or pERV-derived NF-κB (upper panels) and IRF1 (lower panels) motifs within NF-κB and IRF1 ChIP-seq peaks, correspondingly, relative to ETS1 and RUNX1 ChIP-seq peaks. **Left:** TFBS shared with macaque. **Right:** great-ape-specific TFBS. **e, Left:** TFBS identified in consensus sequences of Alu subfamilies that are most abundant in NF-κB ChIP-seq peaks (#>100 copies). The presence of a motif is indicated in black, while its absence is shown in grey. **Middle:** Proportion of enhancer-located Alu elements containing the same TFBS. **Right**: Proportion of Alus containing any of the four NF-κB1 motifs. **f, Left panels:** Difference in binding affinity (Δ) of Alu-derived NF-κB1 motifs (MA00105.4) shared with macaque or chimpanzee, comparing their sequences in the human genome to those in macaque and chimpanzee genomes using TFBStools predictions. **Right panels:** Difference in binding affinity (Δ) of great-ape-specific Alu-derived NF-κB1 motifs compared to their chimpanzee orthologs and the corresponding human consensus sequences. Alu species most abundant in NF-κB ChIP-Seq peaks are pooled to plot cumulative affinity changes.

To evaluate the *in vivo* functionality pTE-derived proinflammatory motifs within enhancers, we analyzed ChIP-seq data from the Remap2022 and ENCODE consortia, covering various tissues. Since NF-κB factors can bind similar motifs through complex and not fully understood interactions^38,39^, we pooled ChIP-seq peaks for NF-κB1, NF-κB2, RELA, and RELB proteins. IRF1 ChIP-seq datasets were also included, as its corresponding motif showed rapid expansion following the human-macaque split. For each pTE biotype, we first quantified the proportions of motifs bound *in vivo* by their corresponding TF (motifs are fully embedded within ChIP-seq peaks). Primate ERVs (pERVs) exhibited the highest proportions of motifs capable of binding, with 40% of NF-κB1 and 24% of IRF1 “shared” motifs found within the corresponding ChIP-Seq peaks (Fig.3b, left panel). In contrast, lower proportions of Alu-derived TFBS are bound, reaching, for example, 8% for “shared” NF-κB1 motifs. This pattern was consistent across both “shared” and “great-ape-specific” motifs (Fig.3b, left and right panels).

We then quantified the proportions of pTE-derived among all motifs bound by either NF-κB or IRF1 within enhancers. Overall, the proportion of pTE-derived motifs was remarkably higher among bound “great-ape-specific” compared to “shared” motifs for both NF-κB and IRF1 (Fig.3c). For instance, over 50% of bound “great-ape-specific” NF-κB1 motifs were pTE-derived, with the majority originating from Alu sequences (Fig.3c, right panel). A similar trend was observed for IRF1 motifs in IRF1 ChIP-seq peaks (Fig.3c). These results indicate that pTE-derived motifs can be functional, and that “great-ape-specific” motifs are intrinsically more prone to binding. Interestingly, Alu-derived motifs were generally less likely to be bound compared to their overall abundance in enhancers (*e.g., OR = 0.14, P-value < 2.2E-16 for “shared” NF-κB1 motifs in NF-κB peaks*). In contrast, pERV-derived motifs were enriched in ChIP-seq peaks relative to their frequency in enhancers (*e.g., OR = 2, P-value < 1.84E-53 for “shared” IRF1 motifs in IRF1 peaks*). Yet, despite lower binding likelihood compared to pERV-derived motifs, Alu-derived ones remained more prevalent within ChIP-seq peaks, both by proportion (Fig.3c) and absolute number of bound motifs (Extended Data Fig.5a), due to their overall abundance. Overall, pERVs overlapped with 4% of IRF1 peaks (Extended Data Fig.5b), while Alus overlapped with 4% to 20% of NF-κB peaks, depending on the specific NF-κB protein (Extended Data Fig.5c).

Given the possibility of accidental overlap with ChIP-Seq peaks, we assessed the specificity of the observed binding. We compared the enrichment of NF-κB motifs (pooled) and IRF1 motifs within their respective ChIP-seq peaks against those of unrelated proteins, such as ETS1 and RUNX1. The analysis revealed significant enrichment of both “shared” and “great-ape-specific” Alu-derived NF-κB motifs in NF-κB peaks, while pERV-derived NF-κB motifs were depleted (Fig.3d, upper panels). Conversely, pERV-derived IRF1 motifs were enriched in IRF1 peaks, whereas Alu-derived IRF1 motifs were depleted (Fig.3d, lower panels). These findings suggest functional specialization, with Alu elements linked to NF-κB activity and pERVs to the IRF1 response.

To determine whether TFBS were present as “ready-to-use” in the original pTE sequences or evolved through random mutations post-insertion, we focused on the Alu and ERV subfamilies most abundant in NF-κB and IRF1 ChIP-seq peaks, respectively. Subfamily consensus sequences from human Dfam^40^ were scanned for the presence of NF-κB and IRF motifs. NF-κB1, NF-κB2, and IRF1 motifs were identified in most Alu subfamilies (Fig.3e, left panel). Furthermore, in enhancer-linked Alus, RELA and RELB motifs, initially absent from consensus, became detectable (Fig.3e, middle panel). While not all Alu elements within enhancers retained the original NF-κB1 or NF-κB2 motifs (Fig.3e, middle panel), the majority contained at least one κB motif type (Fig.3e, right panel). This suggests an evolutionary transition from NF-κB1 and NF-κB2 variants toward highly similar REL motif variants. In contrast to Alus, pERV consensus sequences displayed a broad diversity of IRF motifs, with only MER41B containing NF-κB motifs (Extended Data Fig.6a).

While ChIP-seq data cannot capture all inflammatory triggers, evolutionary shifts in TFBS binding affinity to cognate TF may reflect past selective pressures and, potentially, functional relevance. Using the SearchSeq function in TFBStools^41^, we predicted the alteration of binding affinity for Alu-derived NF-κB1 and pERV-derived IRF1 motifs following the human-macaque split. For “shared” motifs, human sequences were compared to those of macaque and chimpanzee. For “great-ape-specific” TFBS, comparisons were made between human and chimpanzee and between enhancer and subfamily consensus sequences. Nearly half of the “shared” Alu-derived NF-κB1 motifs exhibited higher predicted binding affinity compared to macaque sequences, while a smaller proportion (10%) showed further increases relative to chimpanzee (Fig.3f, left panels). This suggests that the main affinity shift occurred during the evolution of the common human-chimpanzee ancestor. “Great-ape-specific” NF-κB1 motifs showed minimal differences between human and chimpanzee; still, both increased (in AluS) and decreased (in AluY) in affinity compared to consensus sequences from human genome (Fig.3f, right panels), suggesting further human-specific optimizations. Given that AluS elements are more numerous than AluY, the human evolutionary trend appears to favor an overall increase in binding affinity. Similar patterns were observed for the pERV-derived IRF1 motifs (Extended Data Fig.6b).

In summary, pTEs have propagated proinflammatory NF-κB and IRF1 motifs across immune-cell enhancers, becoming particularly rich source of the great-ape-specific binding sites. These motifs increase the activity of enhancers to bind their cognate transcription factors *in vivo* and tend to evolve toward higher binding affinity over evolutionary time.

### Alus are fertile substrates of positive selection in human

To investigate the impact of natural selection on pTEs and their associated enhancers in modern humans, we analyzed three major continental populations from the 1000 Genomes Project (Central Europeans from Utah (CEU), Yoruba from Nigeria (YRI), and Southern Han Chinese (CHS). We assessed positive selection signals in enhancer-overlapping single nucleotide polymorphisms (SNPs) by integrating measures of genetic differentiation between populations (population branch statistics)^42^ with haplotype-based evidence of a rapid increase in derived allele frequency, as provided by Relate^43^. This analysis identified 50,000 enhancer-associated SNPs with signatures of positive selection (PS-SNPs, *Fisher’s combined p-value <0.01*). We further identified positive selection acting on entire enhancer loci by taking the strongest selection *p-values* across enhancer-overlapping SNPs and adjusting for multiple testing to obtain a single selection *p-value* per enhancer. Applying a 5% FDR to these *p-values*, we identified 3,500 enhancers presenting signatures of positive selection of the whole region (PS-enhancers).

We first examined the correlation between pTE presence and positive selection at the enhancer level. Our analysis revealed that enhancers containing Alu elements are significantly more likely to be under positive selection in modern humans than those without, particularly when the Alu sequences were acquired after the split from macaque (*de novo* Alus) (Fig.4a, left panel). In contrast, pERVs and pLINEs did not correlate with positive selection, indicating this effect is Alu-specific. The presence of NF-κB1 motifs further enhances the likelihood of positive selection, especially if these motifs are Alu-derived (Fig.4b). This shows that Alu-derived NF-κB1 motifs alone can reflect specific dynamic properties of enhancers in modern human populations.

**Fig. 4.**
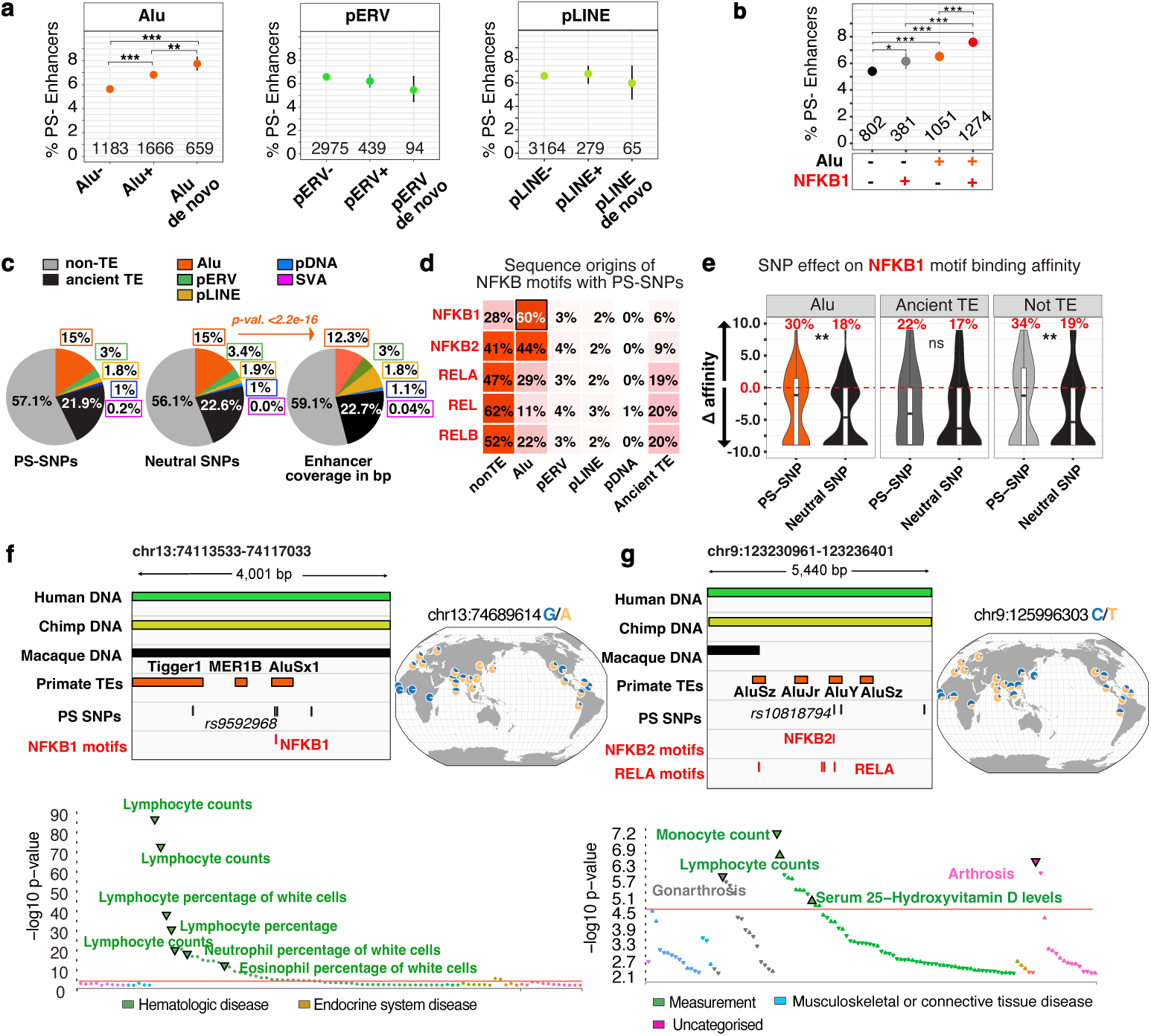
Association of Alus with positive selection in contemporary humans. **a,** Proportion of enhancers under positive selection across different groups: enhancers with and without Alus, pERVs, or pLINEs (shared or absent in the macaque genome, RheMac10). The percentage of positively selected enhancers (5% enhancer-level FDR) in each group is shown. Vertical lines indicate standard deviation across individuals. Numbers of positively selected enhancers are indicated. Fisher’s exact test; *** *P-value 0 - 0.001*, ** *P-value 0.001 - 0.01*. **b,** Proportion of positively selected enhancers either containing Alus or not, and either containing NF-κB-MA0105.4 motif or not (whether Alu-derived or not). **c, Left and middle:** Proportional representation of distinct sequence types harboring PS-SNPs (*p-value < 0.01*) or neutral SNPs (*p-value>0.5*) within enhancers. **Right:** Proportion of the total length of all enhancers covered by different sequence types. Shown is the significance of Alu enrichment within SNP-carrying sequences compared to Alu abundance in enhancer sequences (by bp), calculated using Fisher’s exact test. **d,** Proportional distribution of NF-κB motifs with PS-SNPs based on their sequence origin. **e,** Distribution of NF-κB1-MA0105.4 binding affinity changes (Δ derived allele vs. ancestral) for PS-SNPs and neutral SNPs within enhancers based on sequence origin; in both cases within Alus, ancient TEs or non-TE sequences correspondingly, estimated using *TFBStools*. Fisher’s exact test was applied to the proportion of TFBS with positive Δ binding affinity between PS-SNPs and neutral SNPs in enhancers. *P-value: * <0.05, ** <0.01, ***<0.001*. **f & g,** Genome browser tracks highlighting examples of positively selected PS-SNPs residing in Alu elements, with TFBS highlighted in red. Global selection patterns are visualized using the Geography of Genetic Variants Browser, with PheWas plots generated based on the data from Open Targets Genetics (https://www.genetics.opentargets.org). Shown are traits significantly associated with reported alleles according to the FinnGen, UK Biobank and GWAS Catalog.

Next, we examined how pTEs themselves are targeted by positive selection within enhancers. By annotating enhancer sequences carrying SNPs, we observed that distinct pTE groups, ancient TEs, and non-TE sequences were similarly represented among both PS-SNPs and neutral SNPs (Fig.4c, left and middle panels). This suggests that no specific type of enhancer sequences is favored by positive selection. Instead, Alus emerge as the most abundant pTE target, accounting for 15% of PS-SNPs, largely due to their high frequency. Moreover, the proportion of Alu-derived SNPs is significantly higher than expected based on the overall Alu abundance (15% with PS-SNPs or neutral SNPs compared to 12% of Alu-derived enhancer sequences) (Fig.4c). This implies that Alus are the most mutable elements in enhancers.

To determine whether selection acted directly on pTE-derived NF-κB motifs, we focused on enhancer-linked PS-SNPs overlapping NF-κB motifs. By analyzing the contribution of different enhancer sequence types to NF-κB motifs under positive selection, we found that NF-κB1-MA0105.4 is more frequently associated with Alu elements (60%) than with other sequences (Fig. 4d). Alus also significantly contribute to other related NF-κB motifs under positive selection, whereas other pTEs play a minimal role. At the same time, both PS-SNPs and neutral SNPs occur within Alu-derived NF-κB1 with comparable frequencies, with only a slight increase observed for the most significant PS-SNPs (Extended Data Fig.7a). Furthermore, the overall proportion of SNPs under positive selection within NF-κB1 motifs is similar for Alu-derived and non-Alu-derived sequences (∼6%, Extended Data Fig.7b). Together, this suggests that Alus likely dominate the selection of NF-κB1-MA0105.4 due to being their richest source.

SNPs modify the binding affinity of NF-κB1 motifs, whether originated from Alu elements or less common sources such as ancient TEs or non-TE sequences (Fig.4e). However, PS-SNPs exhibit a distinct pattern compared to neutral SNPs. In Alus, 30% of PS-SNPs increase the binding affinity of NF-κB1 motifs relative to the ancestral allele, compared to only 18% for neutral SNPs (Fig.4e, left panel). Similarly, other positively selected NF-κB motifs show a marked shift toward higher binding affinity compared to neutral SNPs (Extended Data Fig.7c), suggesting that recent positive selection has preferentially favored alleles that increase the affinity of immune enhancers toward NF-κB.

We report two variants with strong selection signals within Alu elements in the PS-enhancers belonging to the intermediate group. The first, rs9592968-A, in the AluSx1-derived NF-κB1 motif shared with macaque (Fig.4f), shows strong selection outside Africa (*p-value_comb.<1E-7, p-value_relate<3E-6, p-value_PBS<3E-3 for CHS*) and increases the binding affinity of this motif (Δ+8.94) relative to the ancestral G allele. The second, rs10818794-T (*p-value_comb<1E-4, p-value_relate<7E-4, p-value_PBS<4E-3 CEU*), reaching highest frequencies in Europe and the Indian Peninsula, resides in the enhancer mainly composed of novel DNA missing from the macaque genome and alters binding affinities of the underlying NF-κB2 and RELA motifs within great-ape-specific AluY (Δ -2.5 and +2.5, respectively) (Fig.4g). Both rs9592968-A and rs10818794-T are significantly associated with altered white blood cell counts (Fig.4f and g, lower panels), potentially linking them to inflammatory and autoimmune conditions^44^.

Our results show that NF-κB binding sites introduced by Alus during primate evolution have served as fertile substrates for adaptation in modern humans. Recent positive selection has modulated the binding affinity of Alu-derived NF-κB1 motifs, likely contributing to the variation in susceptibility to inflammatory disorders among contemporary humans.

### Inflammatory disease-associated enhancers are the most adaptive

To investigate how selection acts upon enhancers associated with inflammatory diseases, which are becoming increasingly prevalent, we constructed a core disease network. We first selected genes from the DisGeNet7.0 associated with major inflammatory and autoimmune diseases and shared by at least one-third of these conditions (Fig.5a). These “core disease genes” were predominantly enriched in myeloid signature and targets for NF-κB1 and IRF1 transcription factors (Extended Data Fig.8a and b). We defined their putative enhancers based on (i) association with these gene promoters according to the ABC contact maps^37^ and (ii) overlap with IRF1 or NF-κB ChIP-seq peaks. This yielded 9,767 regions, referred to hereafter as inflammatory disease enhancers, or IDEs.

**Fig. 5.**
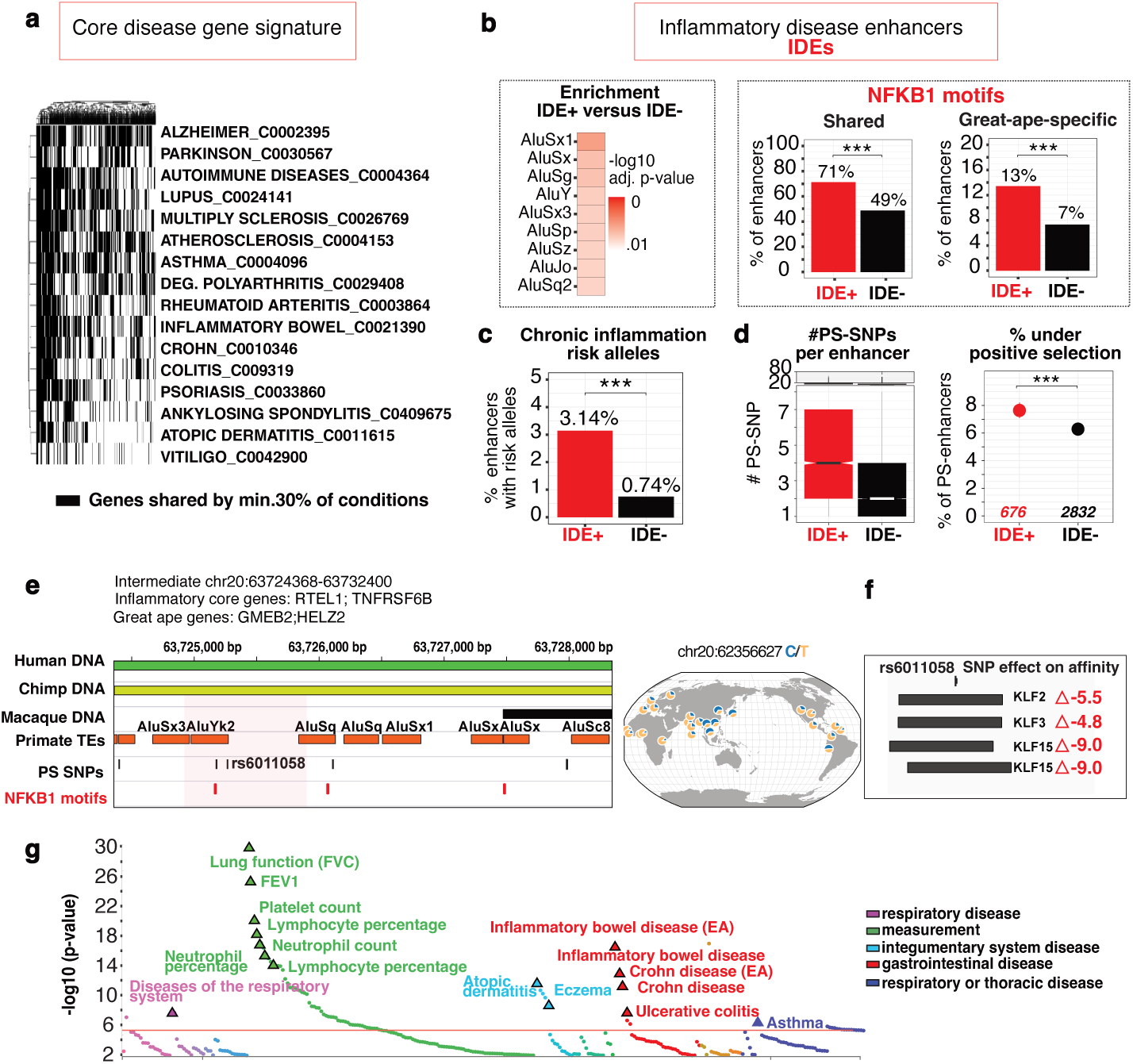
Evolution of the inflammatory disease-associated enhancers. **a,** Binary heatmap displaying genes (in black) shared among at least 30% of the major chronic inflammatory and autoimmune diseases, representing the degree of overlap between different conditions. **b, Left:** Enrichment of Alus in IDEs compared to non-IDEs used as a background. **Right panels:** Proportion of enhancers containing NF-κB-MA0105.4 motifs, whether shared with macaque (left panel) or great-ape-specific (right panel). **c,** Proportion of IDEs and non-IDEs carrying SNPs associated with chronic inflammatory and autoimmune disorders. **d, Left:** Number of PS-SNPs *(P<0.01)* per enhancer within IDE and non-IDE groups. **Right:** Proportion of enhancers under positive selection within IDE and non-IDE groups. Fisher’s exact test; *** *P-value 0 - 0.001.* Numbers of positively selected enhancers are indicated. **e,** Genome browser track highlighting a fragment of the intermediate positively selected IDE carrying PS-SNP rs6011058-T (*P<2E-* 5) residing in the AluYk2 element. In the middle, the global selection patterns are visualized using the Geography of Genetic Variants Browser. **f,** KLF motifs overlapping with rs6011058-T SNP and the binding affinity change of the derived allele versus ancestral accessed using TFBStools. **g,** PheWas plot generated based on the data from Open Targets Genetics (https://www.genetics.opentargets.org) shows traits significantly associated with rs6011058-T according to the FinnGen, UK Biobank and GWAs Catalog.

IDEs demonstrate unique patterns of evolution in primates, being more enriched in Alu elements than other enhancers (set as a background) (Fig.5b, left frame) and more frequently harboring NF-κB1 motifs compared to non-IDEs (71% of IDEs vs. 49% of non-IDEs with “shared” motifs and 13% of IDEs vs. 7% of non-IDEs with “great-ape-specific motifs”) (Fig.5b, right frame). Moreover, IDEs likely play a leading role in the human adaptation of the inflammatory response, with 3% carrying SNPs reported as causative to chronic inflammatory and autoimmune disorders^1^ compared to 0.7% of non-IDEs (Fig.5c). They harbor more PS-SNPs per enhancer overall compared to non-IDEs (Fig.5d, left panel). Furthermore, a higher proportion of IDEs is under current positive selection compared to non-IDEs (Fig.5d, right panel). Together, this suggests that IDEs have been crucial in adapting the inflammatory response throughout primate evolution and remain the most adaptive in modern humans.

We examined IDE association with genes that respond stronger to immune stimulation in great apes compared to monkeys and in humans compared to other primates ^10^. IDEs were strongly associated with both human-chimpanzee-specific and human-specific early transcriptional response genes (Extended data Fig.8c). For instance, over 40% of IDEs were linked to human-chimpanzee-specific specific genes, compared to just 6% of non-IDEs. This suggests a crucial role of IDEs in wiring this evolutionary novel immunity-related transcriptional response in the common human-chimpanzee ancestor.

A compelling example of the recent evolutionary adaptation of IDE is provided by the rs6011058-C/T variant occurring in the great-ape-specific AluYk2 and undergoing strong positive selection in Africa (*p-value_comb < 2E-*5, *p-value-relate < 2E-5, p-value PBS < 1E-2 for YRI*) (Fig. 5E). The surrounding enhancer is primarily composed of great-ape-specific DNA enriched with Alu sequences and harbors an NF-κB1 motif 89 bp away from rs6011058 (Fig.5e). While the derived allele rs6011058-T does not alter an NF-κB1 binding affinity directly, it disrupts predicted binding sites for Krüppel-like factors KLF2, KLF3, and KLF15, (Δ _affinity_ up to -9.0, Fig. 5E). KLFs are known to counteract NF-κB functions and reduce NF-κB-mediated transcriptional activity^45–47^, thus preventing acute and chronic inflammatory conditions^47,48^. rs6011058-T has previously been identified as causal for Crohn’s disease^1^ and is associated with other inflammatory disorders, such as atopic dermatitis and eczema (Fig.5g). The disruption of KLF motifs may thus enhance the effect of NF-κB1 binding, increasing the activity of this IDE in driving expression of its target genes from the core inflammatory signature (RTEL1 and TNFRSF6B), as well as interferon-stimulated, great ape-specific immune response gene HELZ2^49^.

In conclusion, we identified enhancers enriched in Alus and frequently linked to inflammatory diseases. These enhancers exhibit signatures of the inflammatory response adaptation in primates and humans, suggesting their key role in the evolutionary shaping of the human inflammatory response.

## Discussion

Recent research has linked human inflammatory responses to adaptive genetic events in human evolution. Here, we shift the perspective to a deeper evolutionary context, tracing the origins of human immune-cell enhancers to our primate ancestors, whose genomes were profoundly reshaped by the activity of young, primate-specific transposons. We demonstrate that, throughout evolution, these initially parasitic elements seeded enhancers with inflammation-related TFBS, which were subsequently repurposed for binding by their cognate transcription factors. The sharply increased regulatory contribution of pTEs during the evolution of the common human-chimpanzee ancestor likely reflects the rapid phenotypic changes in large anthropoid primates. Such shifts may have required accelerated adaptation, with pTE-derived inflammation-related TFBS providing readily available, albeit potentially opportunistic, genetic tools to fine-tune inflammatory responses. Our findings suggest that pTEs not only shaped the interplay between immunity and environmental pressures in primates but also contributed rich substrates for natural selection and susceptibility to inflammatory diseases in humans.

More specifically, we observed a functional distinction between primate ERVs and Alus. While pERVs appear to be proficient binders of IRF1, as shown here and elsewhere^21^, Alus are specifically biased toward NF-κB binding. Although their binding propensity *in vivo* is relatively low, Alu-derived NF-κB and IRF1 motifs may exert a greater influence on inflammatory responses than those from pERVs, owing to their substantially higher abundance in enhancers. The regulatory significance of Alus has progressively grown throughout evolution, as they became a key source of evolutionary recent, great-ape-specific NF-κB1 motifs. Subsequently, Alus have served as rich substrates for positive selection in humans, potentially contributing to the adaptation of NF-κB responses.

While NF-κB motifs were previously identified in Alu sequences^50,51^, we show that they are embedded in the original Alus, transitioning between similar sequence variants over time and potentially continuously optimizing for better TF binding. Although we show that not all Alu-derived NF-κB motifs are bound by their cognate proteins *in vivo*, their evolutionary fine-tuning may be explained by utilization in response to threats either not replicated in laboratory settings or intermittent usage. It has indeed been shown that κB protein dimers may scan κB sites in a trial-and-error fashion to adjust transcriptional output^39,52^. In this scenario, prevalent Alu-derived NF-κB motifs may have been transiently sampled during proinflammatory stress events and positively selected if proven advantageous. This process may persist today, as the reservoir of unused motifs remains unsaturated yet available, given their presence in the favorable epigenetic environment of enhancers.

We observed that Alus are linked to greater enhancer adaptability in humans and are enriched within the most adaptive enhancers associated with chronic inflammatory diseases. This suggests that Alus play a role in facilitating responses to proinflammatory challenges. This ability may arise from the NF-κB1-MA0105.4 motif, which stands out as evolutionarily significant, having undergone the most rapid expansion in the common human-chimpanzee ancestor and now showing a strong association with adaptive enhancers in humans. Furthermore, as highly mutable elements, Alus may increase evolutionary possibilities and, therefore, the likelihood of enhancers being targeted by natural selection. Finally, Alus are known to favor enhancer-promoter interactions by forming duplexes with complementary sequences^53^ and locally delivering transcription factors^51^. The iterative sampling of promoters by enhancer Alus, transiently bound by NF-κB, could further expand adaptive opportunities. The looping of three distantly located Alus to deliver NF-κB to the IFNb promoter during the early antiviral response, thereby jump-starting gene expression^51^ may serve as an example of such a phenomenon in action. Although we do not explore the broader contribution of Alus to other TFBS, Alu-derived NF-κB antagonists such as NFE2 and KLFs may help to orchestrate a more complex regulation of the inflammatory response.

One might expect the abundance of Alus to dilute the impact of individual elements, with no single Alu being critical. However, depletion of individual copies using CRISPR-Cas9 technology has been shown to disrupt target gene expression^53^. Among these Alus, we found thirty-eight elements whose removal in immune-cell enhancers disrupts the expression of the corresponding target genes.

We propose the concept of regulatory co-option of Alu elements by chance, driven by the law of large numbers. As Alus became prominent in dynamically evolving enhancers, they enabled rapid adaptation to environmental stimuli through pre-existing inflammation-related regulatory motifs. As such, Alu elements may not only reflect the survival history of our primate ancestors but also serve as a vast reservoir for future resilience, potentially ensuring the continued adaptability of the human lineage.

## Acknowledgements

We thank all members of the Amigorena team for constructive feedback. We thank Lluis Quintana-Murci and Camille Berthelot for fruitful comments. Human and other primate icons are borrowed from the Biorender website (https://www.biorender.com). This work was supported by the Program France 2030 launched by the French Government through the LabEx DCBIOL (SA), Agence Nationale de la Recherche, ANR-10-IDEX-0001-02 PSL (SA), Agence Nationale de la Recherche, ANR-23-CE14-0017-02 (EZ), Agence Nationale de la Recherche, grant POPCELL-REG (ANR-22-CE12-0030-01) (MR), MY was supported by Mnemo Therapeutics and the Center of Clinical Investigation, CIC IGR-Curie 1428 of Inserm.

## Author contributions

EZ conceptualized and directed the study. Computational analysis was performed by MY. Population genetics analysis was performed by MR. Data mining and interpretation were performed by EZ, MY, MR. Findings were acquired by SA, EZ, MR. EZ wrote the manuscript. MR and SA edited the manuscript.

