## Supplementary material for "Transposon invasion of primate genomes shaped human inflammatory enhancers and susceptibility to inflammatory diseases": Methods and Extended Data Figures

### Primate-specific transposable elements shape the evolution of inflammation-related enhancers

**The PDF file includes:**

Materials and Methods with references
Figs. S1 to S8

#### Methods

##### Enhancer sequence divergence analysis

Creating the list of putative immune cell enhancers

Coordinates of putative enhancers in human CD34$^+$, CD4$^+$ T cells, CD8$^+$ T cells, B cells, CD19$^+$ cells, monocytes, macrophages, and dendritic cells were obtained from the EnhancerAtlas2.0 database[1] (http://www.enhanceratlas.org/). Coordinates of putative enhancers for human lymphoblastoid cell lines were obtained from *Garcia-Perez et al.*[2]. Enhancer coordinates from EnhancerAtlas 2.0 BED files were converted from the human genome GRCh37 assembly to the human genome GRCh38 assembly using the UCSC *LiftOver* tool[3] with default parameters and the *hg19ToHg38.over.chain* file. Overlapping coordinates were merged, and all enhancers were pooled for further analysis using *BEDtools* (v2.25.0). To select enhancers with accessible chromatin, ATAC-sequencing (ATAC-seq) data for different populations of T cells, B cells, and monocytes, activated or not from *Corces et al.*[4] , was reanalyzed. First, ATAC-seq reads from the same populations, and the same treatment types were merged using *Samtools v1.3* and aligned to the GRCh38 genome using *Bowtie2 v2.2.9 (parameters: -q -N 1 -p 8)*[5]. Peaks were called using *MACS2 v2.1.1 (parameters: -g hs -p 1e-9)*[6], and only robust peaks with a five-fold enrichment over a background were selected. Putative enhancers containing at least one ATAC-peak peak summit were used for further analysis and termed "immune-cell enhancers."

Stratification of human immune-cell genes according to the evolutionary age sequence divergence

To compare the DNA of human immune cell enhancers with sequences from the Rhesus macaque and chimpanzee genomes, enhancer coordinates from human *GRCh38* assembly were converted to the Macaca mulatta (*RheMac10*) and Pan troglodytes (*PanTro6*) genome assemblies using the UCSC *LiftOver* tool with a minimum match threshold of 0.97. Based on the evolutionary age of the species to which enhancers were aligned with 97% similarity, enhancers were categorized as "Static" (full orthologs, human-chimpanzee-macaque alignable), "Intermediate" (human-chimpanzee alignable), or "Rapid" (human-specific variations). Intermediate and rapid enhancers were collectively referred to as 'Dynamic".

Quantification of the mutational rate in enhancer groups since the divergence from macaque

To quantify the mutation rate per enhancer since the divergence from macaque across the three enhancer groups, a comparison between human and macaque DNA was conducted using the *BLASTn* tool (Nucleotide-Nucleotide BLAST 2.13.0+)[7]. First, the genomic origins of dynamic enhancers in macaque DNA were identified by lifting over coordinates from the human *GRCh38* assembly to the *RheMac10* assembly using a minimum match threshold of 0.5 (to obtain quasi-orthologs). Second, a comparison was made between static enhancers and their orthologs in macaques, and dynamic enhancers and their quasi-orthologs, focusing on the percentage of mismatches between human and macaque DNA. To identify evolutionary novel DNA (corresponding to gaps in genomes of chimpanzee and macaque relative to human, we utilized the *UCSC mafNoAlign tool* along with the syntenic alignments from the *hg38.panTro6.synNet* (https://hgdownload.soe.ucsc.edu/goldenPath/hg38/vsPanTro6/) and *hg38.rheMac10.synNet* (https://hgdownload.soe.ucsc.edu/goldenPath/hg38/vsRheMac10/) MAF files.

**Enhancer gene target identification**

The gene-enhancer maps from the activity-by-contact maps (ABC) model [37]. The file *AllPredictions.AvgHiC.ABC0.015.minus150.ForABCPaperV3.txt* was downloaded from https://www.engreitzlab.org/resources/. Enhancer coordinates from immune cell types were lifted to human genome GRCh38 assembly using UCSC *LiftOver* tool with default parameters and the *hg19ToHg38.over.chain* file. If enhancer coordinates from this study overlapped with those from the ABC source by at least 1bp, they were considered to represent the same region. Genes specific to immune response were obtained from the RNA-seq analysis of the human blood PBMC stimulated during 4h with LPS or GARP *ex-vivo*[8].

**TE analysis**

TE subfamily enrichment analysis

To identify subfamilies of transposable elements (TEs) enriched in distinct enhancer groups, genomic coordinates of TEs were retrieved using *RepeatMasker* (*Smit, A.F.A., Hubley, R., & Green, P. RepeatMasker Open-4.0. Repeat Library 20140131* available at http://www.repeatmasker.org). Parsing was performed at both the individual copy and subfamily levels using RepeatMasker output file with the custom *parseRM.pl script*[9]. The ratio of TE subfamily abundance (by copy number and length) within distinct enhancer groups versus the entire genome was analyzed using the `TE-analysis_pipeline.pl` script (https://github.com/4ureliek/Teanalysis). Enrichment significance was assessed using a hypergeometric test with an adjusted *P-value* < *0.01*. To confirm that TE subfamily enrichment was not biased by the length distribution of the number of regions in distinct enhancer groups, we applied the same statistical test to random genomic regions matched to each enhancer group by number and size distribution, excluding regions annotated as enhancers in the *EnhancerAtlas 2.0* database. These random control regions were shuffled 1000 times using *BEDtools* (v2.25.0). Subfamilies uniquely enriched within enhancer regions, with a 5% error tolerance and not enriched in random regions, were defined as "enhancer-enriched TE subfamilies."

TE clade definition

TE clade information was extracted from the file "20141105_hg38_TEage_with-nonTE.txt" downloaded from the TEanalysis tool (https://github.com/4ureliek/Teanalysis).

Trajectory of TE sequence acquisition by enhancer groups over evolutionary time

Genomic gaps in chimpanzee (*PanTro6*) and macaque (*RheMac10*) relative to the human genome *GRCh38* were obtained using the *UCSC mafNoAlign tool* along with the syntenic alignments from the *hg38.panTro6.synNet* (https://hgdownload.soe.ucsc.edu/goldenPath/hg38/vsPanTro6/) and *hg38.rheMac10.synNet* (https://hgdownload.soe.ucsc.edu/goldenPath/hg38/vsRheMac10/) MAF files.

Coordinates of primate-specific transposable elements (pTEs) were overlapped with these gaps using the *BEDtools intersectBed tool* (-f 0.9). If a primate-specific TE (pTE) sequence was present in the same genomic location in macaque, it was classified as shared. If the sequence mapped to a genomic gap in macaque or chimpanzee, regardless of the mapped length, it was considered great-ape-specific or human-specific, respectively. For Fig.1f, a TE copy was classified as a new insertion if it overlapped with the gap by at least 90% of its length.

pTEs in ATAC-seq peaks

ATAC-seq peaks from distinct immune-cell populations were classified as "shared" if more than 50% of their length aligned with the macaque genome *(RheMac10)*. If more than 50% of an ATAC-seq peak matched genomic gaps in *RheMac10*, it was classified as "novel." ATAC-seq peaks were considered Alu-derived or pERV-derived if at least 50% of their sequence consisted of Alu or pERV elements. Among the novel ATAC-seq peaks, those where over 50% of the gap overlapped with Alu or pERV sequences were defined as "novel Alu- or pERV-derived peaks." The proportion of Alu- or pERV-derived peaks, whether novel or shared, was quantified relative to the total number of novel or shared peaks, respectively.

Proportion of the total length of all enhancers covered by sequences of different types

For Fig.4c, RepeatMasker output was customized using TE clade information from https://github.com/4ureliek/Teanalysis classifying TEs into primate-specific and ancient. Primate-specific TEs were further classified according to biotype (Alus, pERVs, pLINA, pDNA, SVA). This file was processed using the parseRM.pl script (https://github.com/4ureliek/Teanalysis) to generate coverage of each primate group and ancient TEs in genome. The output file,

<RMout.out>.parseRM.all-repeats.tab, reported the genomic coverage for each group. Overlapping regions between different groups accounted for 0.02% of the total and were excluded from the analysis. Coverage in enhancers was further calculated using TE-analysis_pipeline.pl script (-TEov 1) (https://github.com/4ureliek/Teanalysis), with input from the parseRM.pl outputs and the same customized file. Coverage of enhancers by nonTE sequences were defined as 100%-SUM(TE-coverage).

**Transcription factor binding site analysis**

TFBS enrichment analysis

TFBS enrichment analysis across the three individual enhancer groups was performed using the Homer (v4.11) *"findMotifsGenome.pl"* tool with the parameter *"-size given"*, utilizing a custom Homer TFBS database (http://homer.ucsd.edu/homer/motif/motifDatabase.html) and a significance threshold of *P<0.01*. To compare the enrichment or depletion of inflammation-related transcription factor (TF) motifs between groups, genomic localizations of IRF and NF-κB family proteins were retrieved from the human JASPAR2022 Transcription Factors Tracks. (http://expdata.cmmt.ubc.ca/JASPAR/downloads/UCSC_tracks/2022/hg38/). A two-tailed hypergeometric test was applied to assess the enrichment of TF motif length coverage in one enhancer group, using another enhancer group as a background. TFBS were considered significantly enriched or depleted with an adjusted *P-value* threshold of 10^-10.

Identifying TFBS shared with macaque or great-ape-specific

Great-ape- or human-specific TFBS were defined by 100% overlap with gaps in the macaque (*RheMac10*) or chimpanzee (*PanTro6*) genomes, respectively, compared to the human genome *GRCh38*. Gaps were identified as above, using *UCSC mafNoAlign tool* along with the syntenic alignments from the hg38.panTro6.synNet (https://hgdownload.soe.ucsc.edu/goldenPath/hg38/vsPanTro6/) and hg38.rheMac10.synNet (https://hgdownload.soe.ucsc.edu/goldenPath/hg38/vsRheMac10/) MAF files.

To access the proportion of Alu elements carrying these TFBS within enhancers, enhancer-linked Alus containing full-length TFBS were identified using *BEDtools intersectBed* (-F 1) and divided by the total number of Alu elements overlapping with enhancers.

Identifying pTE-derived TFBS shared with macaque or great-ape-specific in enhancers

To investigate the contribution of primate-specific transposable elements (pTEs) to the creation of TFBS within enhancers, TFBS coordinates were obtained from the human JASPAR2022 Transcription Factors Tracks (as above). TFBS were classified as either "shared" (present at the same genomic location in macaques (*RheMac10*) or "great-ape-specific" (mapping entirely to a gap in the macaque genome (*RheMac10*) compared to the human genome (*GRCh38*), with gaps identified as described above). To assess the contribution of pTEs, TFBS were overlapped 100% with pTE sequences, and pTEs were grouped by biotype (Alu, primate-specific ERVs, LINEs, DNA elements [pERV, pLINE, pDNA], and SVA). The proportion of TFBS derived from each pTE biotype was then calculated separately for shared and great-ape-specific TFBS motifs, relative to the total number of each TFBS type in enhancers.

Analysis of TFBS in ChIP-seq peaks within enhancers

ChIP-seq peak coordinates for IRF and NF-κB family proteins were collected from ENCODE (https://www.encodeproject.org) and REMAP2022 database[10] (https://doi.org/10.1093/nar/gkab996) and pooled at TF family level (with NF-κB1, NF-κB2, RELA and RELB combined together) using *BEDtools* with default parameters. To identify NF-κB motifs (NF-κB1_MA0105.4, NF-κB2_MA0778.1, RelA_MA0107.1, and RelB_MA1117.1) and IRF1_MA0050.2 motifs in enhancer ChIP-seq peaks, coordinates of these motifs the human JASPAR2022 Transcription Factors Track (as above) were overlapped with coordinates of their corresponding ChIP-seq peaks using *Bedtools*. Only motifs that overlapped 100% with ChiP-seq peaks were selected as bound TFBS. To quantify the proportion of bound TFBS within each pTE group, the ratio of bound motifs to the total number of corresponding motifs in enhancers was calculated. To assess the proportional distribution of bound TFBS (shared or great-ape-specific) based on their sequence origin (pTE biotypes and non-TE), we quantified the proportion of bound motifs assigned to distinct pTE biotypes or the non-TE group relative to all bound TFBS of the same type.

For the proportion of ChIP-seq peaks overlapping with Alu or pERV sequences, pTEs were considered overlapping if at least 6bp of the pTE sequence intersected with a peak.

To study the enrichment or depletion of Alu-derived NF-κB motifs within NF-κB ChIP-seq peaks or IRF1 motifs within IRF1 peaks, an arbitrary background was created using ChIP-seq peaks from REMAP2022, uniquely attributed to unrelated ETS1 and RUNX1 proteins (not overlapping with NF-κB or IRF1 ChIP-seq peaks). A Fisher's exact test was then conducted to compare the ratio of Alu-derived or pERV-derived NF-κB motifs (NF-κB1_MA0105.4, NF-κB2_MA0778.1, RELA_MA0107.1, and RELB_MA1117.1 pooled together) to non-Alu-derived or non-pERV-derived NF-κB motifs within NF-κB ChIP-seq peaks against their ratio within ETS1 or RUNX1 ChIP-seq exclusive peaks, to identify significantly enriched or depleted motifs (*P-value < 0.01*). The same statistical analysis was applied to Alu-derived or pERV-derived IRF1 motifs in IRF1 ChIP-seq peaks.

#### Identifying TFBS in pTE consensus sequences

To identify IRF and NF-κB family protein motifs in the progenitors of enhancer-linked pTE copies (for subfamilies of Alus and pERVs), their subfamily consensus sequences were extracted from Dfam human collection of consensus models (https://dfam.org)[11 40] using *Dfam.embl* file and converting to *FASTA* format using web-based "*embl_to_fasta*" tool (https://sequenceconversion.bugaco.com/converter/biology/sequences/embl_to_fasta.php). To measure the significance of the candidate TFBS in matching pTE subfamily *Dfam* consensus sequences, *FIMO* tool (https://meme-suite.org/meme/tools/fimo) was utilized with default parameters using TFBS matrix from JASPAR2022.

#### In silico prediction of enhancer-resident TFBS binding affinity

Nucleotide sequences of TFBS that overlapped 100% with enhancer-linked Alus or pERVs were retrieved using the *Biostrings::getSeq* function in R (version 4.2.2) and the BSgenome object *Hsapiens.UCSC.hg38* (R package version 1.4.5). These TFBS were lifted to the chimpanzee (*PanTro6*) and macaque (*RheMac10*) genome assemblies using the UCSC *LiftOver* tool with a *-minMatch parameter of 0.5* to obtain motifs shared between human and other primates. Shared TFBS nucleotide sequences for macaque and chimpanzee were extracted using the *Biostrings::getSeq* function with the *BSgenome.Ptroglodytes.UCSC.panTro6* and *BSgenome.Mmulatta.UCSC.rheMac10* objects. TFBS binding affinity was quantified using the

*TFBSTools::searchSeq* function in R[12]. To assess the evolution of TFBS binding affinity for Alu-derived NF-κB motifs and pERV-derived IRF1 motifs since the human-macaque divergence, affinities of shared TFBS were compared between human and macaque or chimpanzee. For great-ape-specific TFBS (absent in the macaque genome), comparisons were made between human and chimpanzee sequences, as well as between human TFBS in enhancers and subfamily consensus sequences. The difference (Δ) in TFBS binding affinity was calculated by comparing maximum scores generated by *TFBSTools::searchSeq* in R. The trajectory of TFBS binding affinity was classified as increasing if $\Delta \geq 2$, decreasing if $\Delta \leq -2$, and neutral otherwise.

**Positive selection in modern human populations**

Positive selection at immune-cell human enhancers

To estimate how immune enhancers have been targets of positive selection in recent human history, we focused on Central Europeans from Utah (CEU), Yoruba from Ibadan, Nigeria (YRI), and Southern Han Chinese from Hong Kong (CHS) populations from the 1000 Genomes Project, representing European, African, and East Asian ancestries, respectively. For each population, allele ages and frequency data, as well as p-values indicating evidence of positive selection calculated using Relate, were downloaded from Zenodo. (https://zenodo.org/records/3234689).

We then measured evidence of positive selection at each variant by combining two orthogonal metrics:

(i) The Relate *p-value* for positive selection[13] that contrasts the age of the derived allele (as inferred from local haplotypic patterns) to that of genome-wide SNPs matched for allele frequency, allowing to detect rapid increases in derived allele frequency.

(ii) An empirical $p$-value derived from the population branch statistic ($PBS$), that captures population-specific changes in allele frequency[14]. For each population, PBS was computed based on Reynold's FST estimates[15], using the other two populations as control and outgroup. For each population, genome-wide PBS values were then ranked across all common variants (minor allele frequency >5%), and each variant $v$ was assigned an empirical $p$-value $p_v$ defined as the percentage of common variants with a $PBS$ value greater than $PBS(v)$.

For each variant and population, the two p-values were then combined using Fisher's method to obtain a combined $p$-value of positive selection. For each enhancer, we next applied Sidak's multiple testing correction across all variants overlapping the enhancer and all three populations and focused on the variant and population with the lowest adjusted *p*-value to obtain a single *p*-value of positive selection per enhancer. Finally, we applied Benjamini-Hochberg (BH) multiple testing correction across all enhancers and applied a 5% FDR threshold to define the set of enhancers evolving under positive selection.

Differences in the percentage of selected enhancers across groups of enhancers were tested using Fisher's exact test. For each group, we derived the 95% confidence interval of the percentage of enhancers under positive selection using the *binom.test* R function.

SNP effect on TFBS binding affinity

Positively selected SNPs (PS-SNP) within immune-cell enhancers were defined by *P-value* of positive selection < *0.01*. Neutrally evolving SNP counterparts were defined as SNP with *p-value >0.5* also located within immune-cell enhancers. Binding affinity comparisons between derived and ancestral alleles was performed using *TFBSTools::searchSeq* R function as above.

**Identification and characterization of inflammatory disease enhancers (IDEs)**

Genes associated with the main inflammatory and autoimmune disorders were downloaded from the DisGenNet database (https://www.disgenet.com), and those overlapping with at least 30% of the conditions were selected. Activity-by-Contact maps *(37)* were used to link these genes to their putative enhancers. Coordinates of these enhancers were intersected with the coordinates of ChIP-seq peaks for IRF1, NF-κB1, NF-κB2, RELA, and RELB from ReMAP 2022 and ENCODE using *BEDtools* and enhancers containing at least 50% of ChIP-seq peaks length (no matter what TF) were designated as inflammatory disease enhancers (IDEs). To identify TE subfamilies enriched in IDE compared to other enhancers, the TE subfamily enrichment analysis described above was conducted. To identify proportion of IDEs associated with genes overexpressing during early immune response in great apes or human, these genes were extracted from[8].

**Data visualization**

Data was visualized in the genome browser IGV 2.8.13.

Global selection patterns were visualized using the Geography of Genetic Variants Browser (https://popgen.uchicago.edu/ggv/?data=%221000genomes%22&chr=1&pos=222087833), PheWas plots were created using data from Open Targets Genetics site, choosing association of traits from FinnGen, UK Biobank, and GWAS Catalog (https://www.genetics.opentargets.org).

**R session information**

R version 4.2.2 (2022-10-31)

Platform: x86_64-conda-linux-gnu (64-bit)

Running under: Ubuntu 16.04.7 LTS

Attached packages:

BiocManager_1.30.23, GenomicRanges_1.50.2, GenomeInfoDb_1.34.9, IRanges_2.32.0, BiocGenerics_0.44.0, Biostrings_2.66.0, S4Vectors_0.36.2, TFBSTools_1.36.0, BSgenome_1.66.3, BSgenome.Hsapiens.UCSC.hg38_1.4.5, BSgenome.Ptroglodytes.UCSC.panTro6_1.4.2, BSgenome.Mmulatta.UCSC.rheMac10_1.4.2, dplyr_1.1.4, ggplot2_3.5.1, ComplexHeatmap_2.14.0.

**Data availability**

Additional information is available upon request.

**Methods references**


1. Gao, T. & Qian, J. EnhancerAtlas 2.0: an updated resource with enhancer annotation in 586 tissue/cell types across nine species. *Nucleic Acids Research* gkz980 (2019)
2. García-Pérez, R. *et al.* Epigenomic profiling of primate lymphoblastoid cell lines reveals the evolutionary patterns of epigenetic activities in gene regulatory architectures. *Nat Commun* **12**, 3116 (2021).
3. Kent, W. J. *et al.* The human genome browser at UCSC. *Genome Res* **12**, 996–1006 (2002).
4. Langmead, B. & Salzberg, S. L. Fast gapped-read alignment with Bowtie 2. *Nat Methods* **9**, 357–359 (2012).
5. Zhang, Y. *et al.* Model-based analysis of ChIP-Seq (MACS). *Genome Biol* **9**, R137 (2008).
6. McGinnis, S. & Madden, T. L. BLAST: at the core of a powerful and diverse set of sequence analysis tools. *Nucleic Acids Res* **32**, W20-25 (2004).
7. Nasser, J. *et al.* Genome-wide enhancer maps link risk variants to disease genes. *Nature* **593**, 238–243 (2021).
8. Hawash, M. B. F. *et al.* Primate innate immune responses to bacterial and viral pathogens reveals an evolutionary trade-off between strength and specificity. *Proc. Natl. Acad. Sci. U.S.A.* **118**, e2015855118 (2021).

#### Extended Data Figures

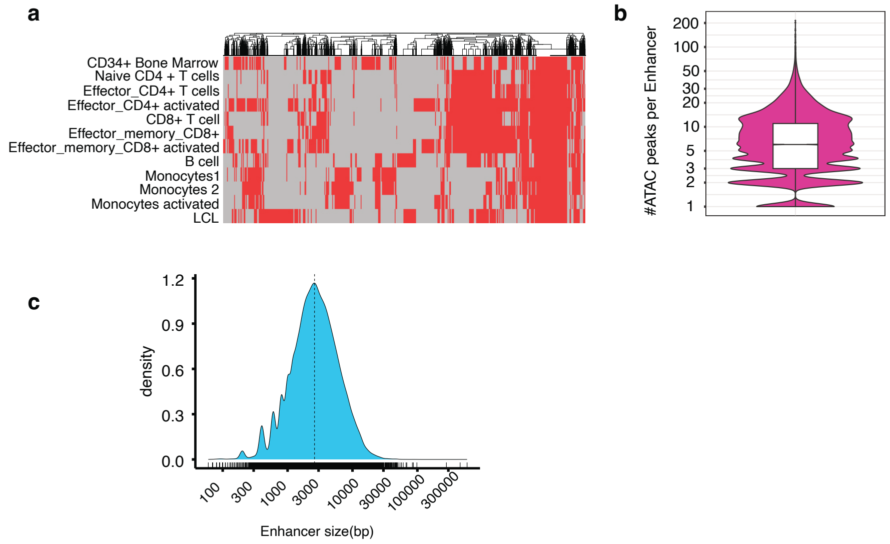


**Extended Data Figure 1 | Characterization of immune-cell enhancers. a,** Binary map demonstrates a degree of enhancer share between immune cell types and represents the presence (red) or absence (grey) of regions annotated as enhancers in immune cells **b,** Number of non-redundant ATAC-peaks pooled from different immune cell populations per enhancer. **c,** Distribution of the enhancer size in bp.

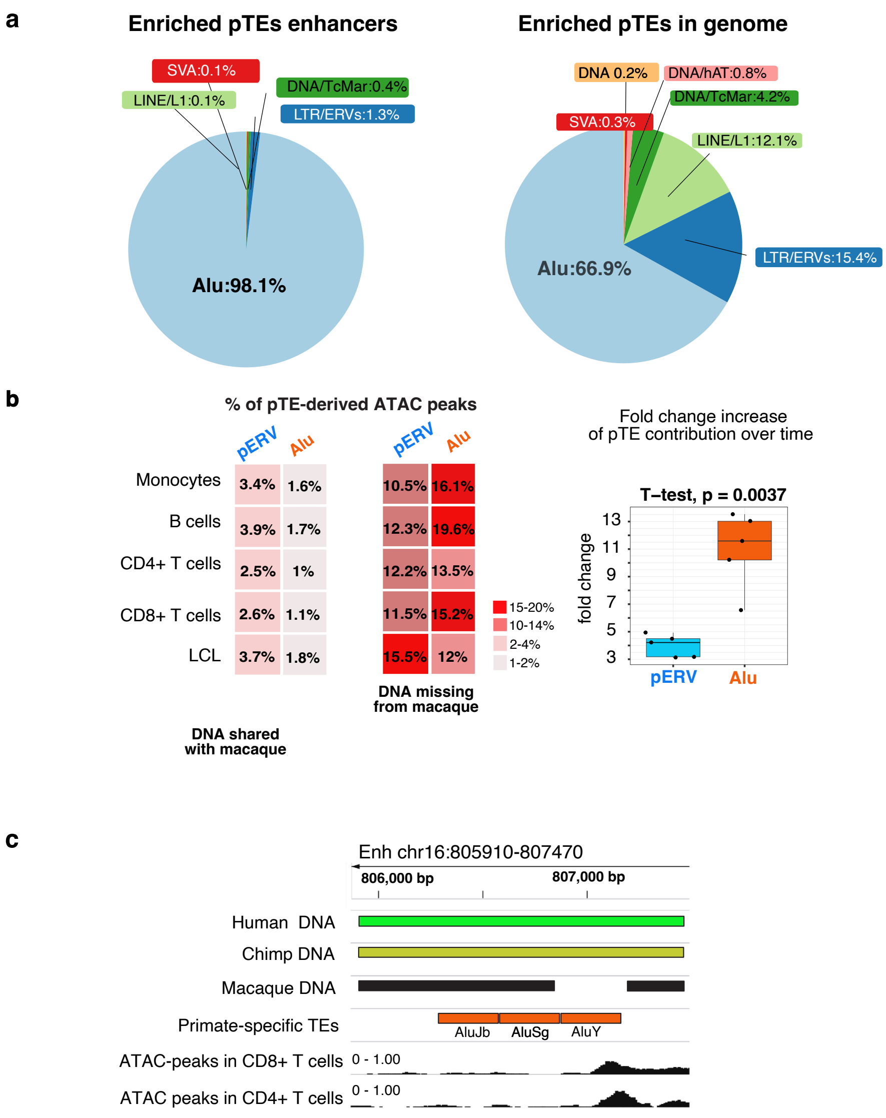


**Extended Data Figure 2 | Alu elements in enhancers. a,** Proportions of primate-specific TEs biotypes of the total enhancer-enriched pTE copies in enhancers and in genome. Subfamilies are grouped according to phylogeny. **b,** Proportion of immune-cell ATAC-seq peaks, mapping (shared) or not (novel) by at least 50% to the RheMac10 and derived from either primate-specific ERV or Alu elements. **Right panel:** Fold change increase in the proportion of ERV or Alu-derived ATAC-seq peaks overlapping with great-ape-specific DNA compared to DNA shared with macaque. **c,** Genome browser track provides an example of the enhancer expanded via the AluY element after the split from macaques and displaying open chromatin (ATAC-seq reads) in human immune cells.

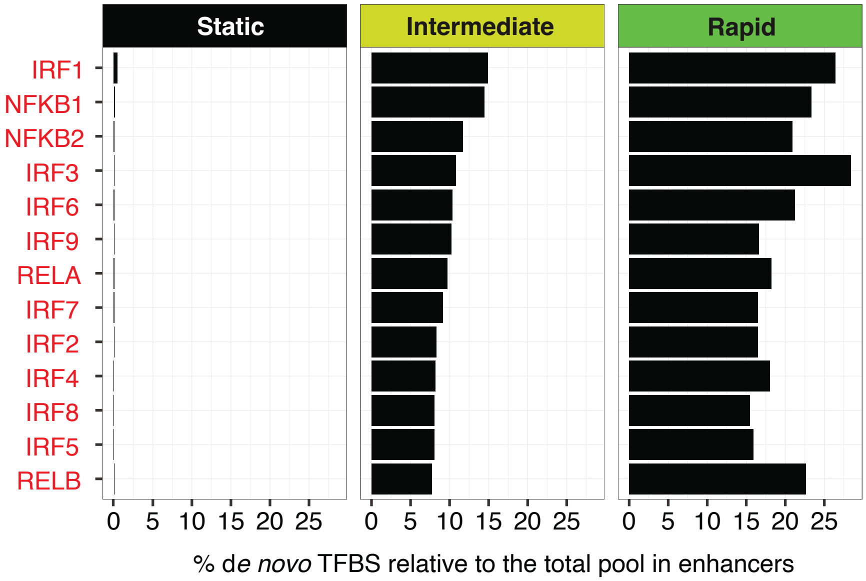


**Extended Data Figure 3 | Proportion of putative TFBS gained since the divergence from macaque in distinct enhancer groups.** Proportion of TFBS corresponding to the DNA in human absent (matching genomic gaps) in the macaque genome from total TFBS of the same kind in each enhancer category.

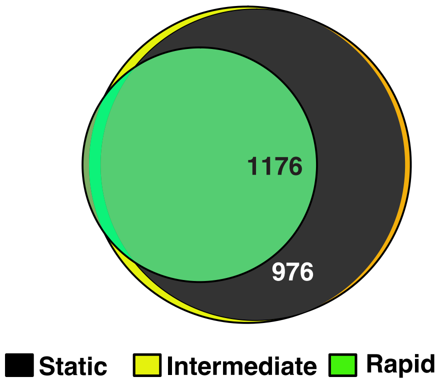


**Extended Data Figure 4 │Overlap between gene targets of distinct enhancer groups.** Overlap between genes assigned to distinct enhancer group according to the activity-by-contact (ABC) enhancer-gene interaction maps. Only genes upregulated after 4h stimulation of human blood cells by infectious agents from *Hawash et al. (ref 9 in the main text)* are shown.

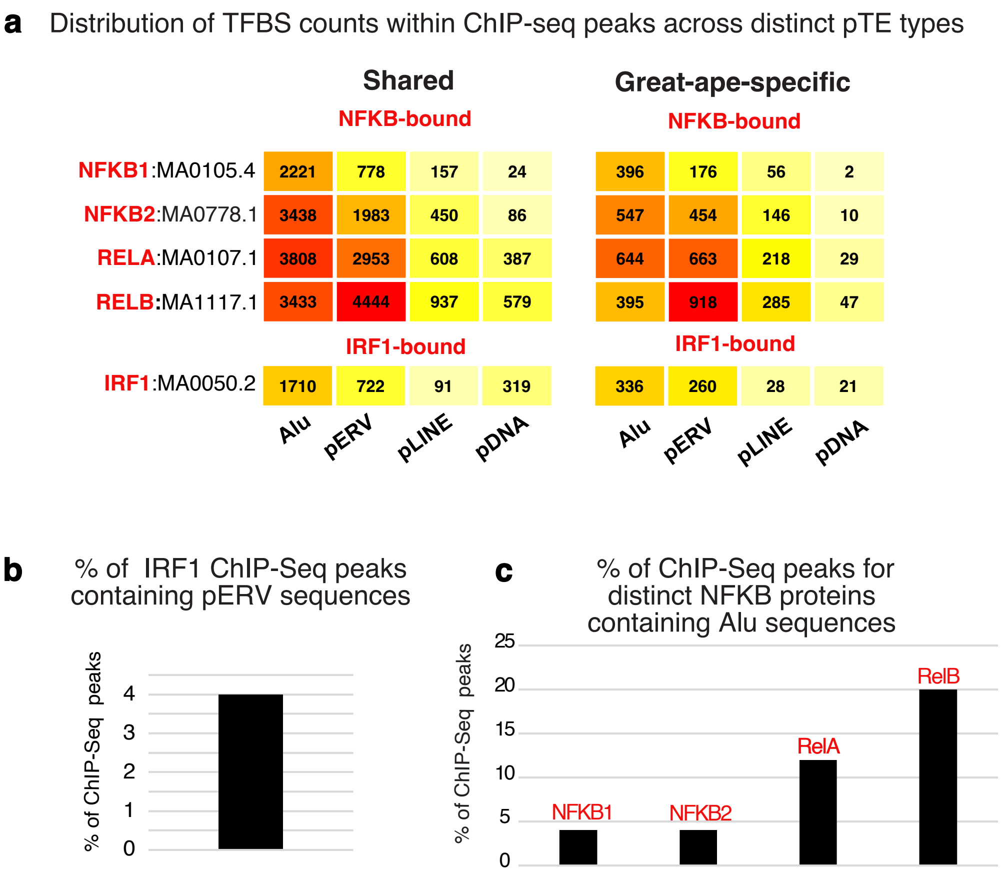


**Extended Data Figure 5 | TFBS and pTEs in ChIP-seq peaks. a,** Number of NF-κB motifs within NF-κB ChIP-seq peaks and IRF1 motifs within IRF1 ChIP-seq peaks, categorized by distinct pTE biotypes. **b,** Proportion IRF1 ChIP-seq peaks overlapping pERVs**. c,** Proportion of ChIP-seq peaks for the distinct NF-κB proteins overlapping with Alu elements.

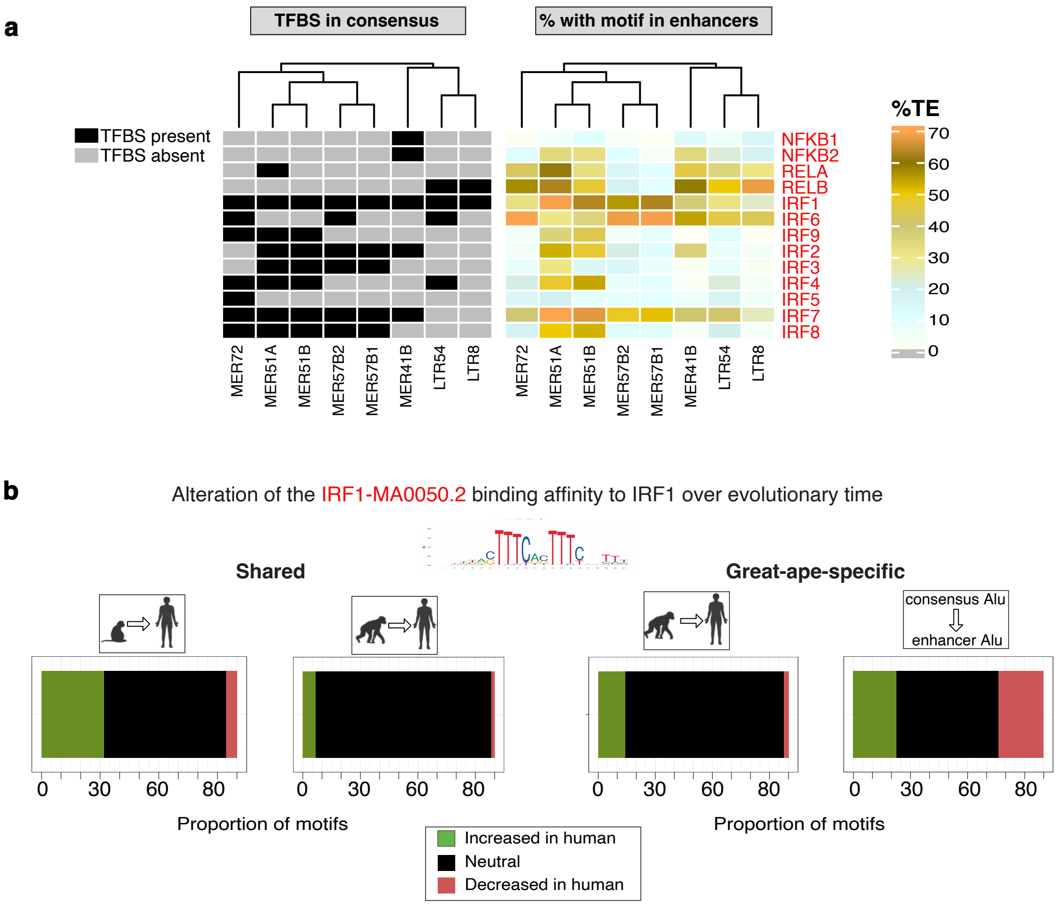


**Extended Data Figure 6 | Evolution of pERV-derived IRF motifs. a, Left panel**: TFBS identified in consensus sequences of primate-specific ERV (pERV) subfamilies that are most abundant (#≥10) within IRF1 ChIP-seq peaks. Presence of a motif is marked in black, and its absence in grey. **Right panel:** Proportion of enhancer-resident pERVs containing the same TFBS. **b, Left panels:** Change in binding affinity (Δ) of IRF1 motifs in human enhancer-resident pERVs compared to their orthologs in macaque and chimpanzee genomes, predicted using TFBStools. **Right panels:** Change in binding affinity (Δ) of great-ape-specific pERV-derived IRF1 motifs compared to their chimpanzee orthologs and the corresponding subfamily consensus sequences. pERV species most abundant in IRF1 peaks are pooled to plot cumulative affinity changes.

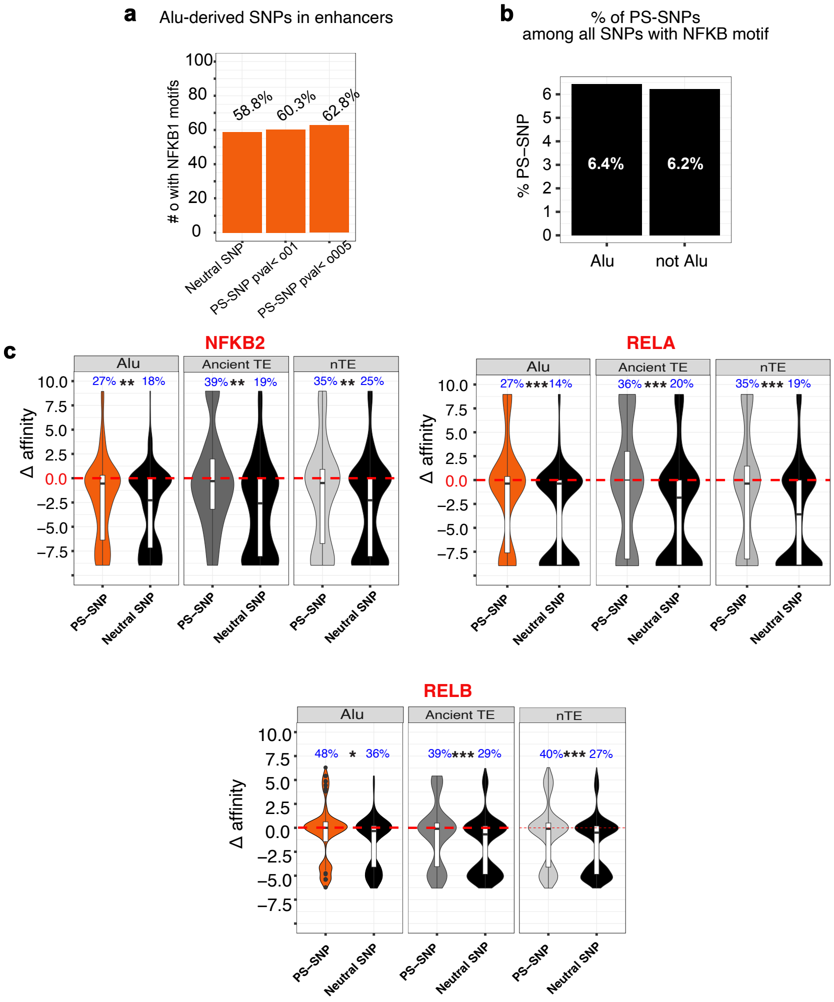


**Extended Data Figure 7 │Characterization of Alus under current positive selection. a,** Proportion of Alu-derived NF-κB-MA0105.4 motifs overlapping with enhancer PS-SNPs across different significance ranges, as well as neutral SNPs (*p-value>0.5*), frequency-matched in the same population, relative to the total number of NF-κB1-MA0105.4 motifs in each category. **b,** Proportion of PS-SNPs among all SNPs with the NF-κB1-MA0105.4 motif, Alu-derived or non-TE-derived. **c,** Distribution of NF-κB motif binding affinity changes (Δ derived allele vs. ancestral) for PS-SNPs and neutral SNPs, both located in enhancers, based on sequence origin, with ancient TEs and non-TE sequences as secondary NF-κB1 motif contributors, estimated using *TFBStools*. Fisher's exact test was applied to the proportion of TFBS with positive Δ binding affinity between PS-SNPs and neutral SNPs in enhancers. *P-value: * <0.05, ** <0.01, ***<0.001.*

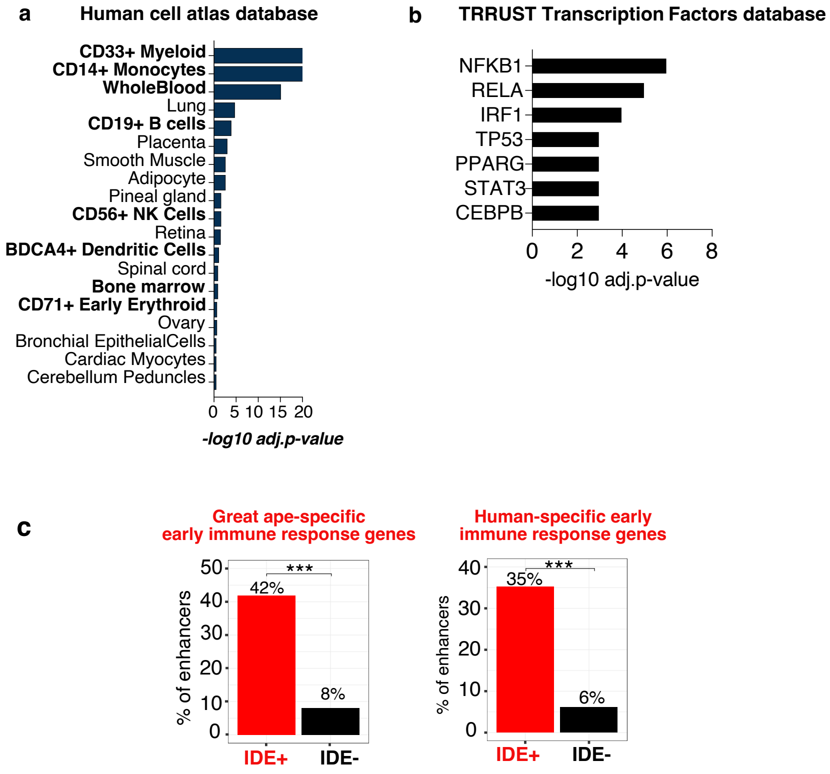


**Extended Data Figure 8 | Characterization of enhancers associated with inflammatory disease. a,** Enrichment of genes linked to at least 30% of autoimmune disorders across different tissues (signature of inflammation), identified using the Human Cell Atlas tool in the EnrichR hub. **b,** Signature of inflammation enriched in transcription factor targets, accessed using the TRRUST Transcription Factors 2019 tool in the EnrichR hub. **c,** Proportion of enhancers in physical contact with genes that show a stronger early transcriptional immune response in great apes (including humans) compared to macaques and in humans compared to great apes, as determined by ABC maps. Gene information is obtained from *Hawash et al.*[10]. Shown is *P-value* with respect to non-IDE enhancers. *** $P \leq 0.001$.
